# Functional Topography of the Neocortex Predicts Covariation in Complex Cognitive and Basic Motor Abilities

**DOI:** 10.1101/2023.01.09.523297

**Authors:** Ethan T. Whitman, Annchen R. Knodt, Maxwell L. Elliott, Wickliffe C. Abraham, Kirsten Cheyne, Sean Hogan, David Ireland, Ross Keenan, Joan H. Lueng, Tracy R. Melzer, Richie Poulton, Suzanne C. Purdy, Sandhya Ramrakha, Peter R. Thorne, Avshalom Caspi, Terrie E. Moffitt, Ahmad R. Hariri

## Abstract

Although higher-order cognitive and lower-order sensorimotor abilities are generally regarded as distinct and studied separately, there is evidence that they not only covary but also that this covariation increases across the lifespan. This pattern has been leveraged in clinical settings where a simple assessment of sensory or motor ability (e.g., hearing, gait speed) can forecast age-related cognitive decline and risk for dementia. However, the brain mechanisms underlying cognitive, sensory, and motor covariation are largely unknown. Here, we examined whether such covariation in midlife reflects variability in common versus distinct neocortical networks using individualized maps of functional topography derived from BOLD fMRI data collected in 769 45-year old members of a population-representative cohort. Analyses revealed that variability in basic motor but not hearing ability reflected individual differences in the functional topography of neocortical networks typically supporting cognitive ability. These patterns suggest that covariation in motor and cognitive abilities in midlife reflects convergence of function in higher-order neocortical networks and that gait speed may not be simply a measure of physical function but rather an integrative index of nervous system health.

## INTRODUCTION

Higher-order cognitive and lower-order sensorimotor abilities have been generally regarded as distinct and studied separately. However, there is growing evidence that they not only covary but also that this covariation increases across the lifespan. For example, there is some evidence for covariation between sensorimotor and cognitive abilities during childhood (van der Fels et al., 2016; Welch & Dawes et al., 2007), midlife (Rasmussen et al., 2019), and old age (van der Wilik et al., 2021). In addition, this covariation tends to become more pronounced during aging (Baltes & Lindenberger 1997). Moreover, measures of sensory and motor abilities such as hearing tests and gait assessments are commonly used to predict cognitive decline and dementia risk in older adults (Collyer et al., 2022; Kwok et al., 2022; Studenski et al., 2011). Identifying brain mechanisms linking these seemingly disparate abilities can further our understanding of not only the extent to which variability in fundamental aspects of human behavior reflect distinct and common features of brain function across the lifespan but also how they may be best leveraged in clinical applications.

Gait speed is a simple and widely used measure of biological aging measured by how quickly a person walks across a biosensor-equipped pad (Fritz & Lusardi, 2009). As early as midlife, individual differences in gait speed covary with complex psychological functions including memory and cognitive ability (Rasmussen et al., 2019), yet the extent to which these processes are associated with common brain systems is unclear. There is some evidence in older adults that gait speed is associated with the strength of functional connectivity in higher-order brain networks supporting cognitive abilities (Lo et al., 2017; Yuan et al., 2015), bolstering the hypothesis that some effects of physical and cognitive aging reflect shared features of common brain systems. Likewise, hearing ability as measured by pure tone audiometry is associated with cognitive ability during midlife and aging (Loughrey et al., 2018; Okley et al., 2021; Lindenberger & Baltes 1994; Baltes & Lindenberger 1997) and is predictive of future cognitive decline (Lin et al., 2011). However, there is some added complexity in assessing potential links between hearing and cognitive ability.

For example, pure tone audiometry is not a consistent indicator of functional hearing ability and some individuals who have no peripheral damage to the ear (i.e., peripheral hearing loss) may still report reduced hearing ability (Sardone et al. 2019), which could be more indicative of underlying brain health. Conversely, peripheral synaptic pathologies have been associated with normal pure tone thresholds (Kohrman et al., 2020). For this reason, hearing researchers often measure declining ability to differentiate speech in noisy environments, which is thought to be more affected by brain processing and is also associated with cognitive decline (Humes et al., 2013, 2020; Sardone et al., 2019; Stevenson et al., 2022).

This ability to recognize speech in the presence of background or competing noise is measured as a speech reception threshold in noise (henceforth “SRT-hearing”). Crucially, measured SRT-hearing may show greater ecological validity than pure tone audiometry, as it captures ability to hear and discern complex auditory information (Stevenson et al., 2022). As with gait speed, prior studies have associated SRT-hearing with functional connectivity in higher-order brain networks most associated with complex cognitive ability (Fitzhugh, et al., 2021). These prior findings with gait speed and SRT-hearing suggest that physical and cognitive aging are reflected in shared features of common brain networks. However, they are limited by small sample sizes and only focus on a small number of networks. Thus, it remains unclear why basic sensorimotor functioning is so closely linked with cognition functioning especially in midlife and continuing into later life.

Intrinsic functional connectivity is a powerful tool to describe individual differences in network-level brain organization and their relationship to behavior (Chen et al., 2022; Shen et al., 2017). Traditional studies of functional connectivity use group-averaged atlases to assign anatomical brain regions to different functional networks. However, this assumes that the spatial layout of functional networks is identical from person to person. Recent technical advances have demonstrated that there is substantial variation between people in the spatial layout of functional neocortical networks (Laumann et al. 2015; Wang et al. 2015). This spatial variation is called *functional topography*. Traditional studies using functional connectivity ignore variation in functional topography, and this likely increases error in the calculation of functional connectivity. Importantly, emerging studies have demonstrated that functional topography reliably maps onto individual differences in behavior as well as onto developmental changes in early life (Kong et al. 2019; Cui et al. 2020, 2022; Keller et al., 2022). Thus, functional topography represents a novel measure that can reduce individual-level error in the estimation of neocortical network architecture and, subsequently, further capture covariation between brain and behavior.

Here, we leveraged measures of functional topography in a large population-representative birth cohort, the Dunedin Study, to examine the extent to which overlap in cognitive, motor, and sensory abilities reflect shared variability in neocortical network architecture. Specifically, we used Multi-Session Hierarchical Bayesian Modeling (Kong et al., 2019; MS-HBM) to generate reliable individualized estimates of functional topography at age 45. We then mapped functional topography onto variation in cognitive ability as measured by IQ, motor ability as measured by gait speed, and sensory ability as measured by SRT-hearing. Next, we tested our ability to predict variation in these three behaviors in Study members by training models on features of functional topography. Based on prior work, we hypothesized that IQ, gait speed, and SRT-hearing would be correlated within each Study member. We further hypothesized that higher IQ, faster gait speed, and better SRT-hearing would all be associated with relatively larger higher-order functional neocortical networks. Lastly, we hypothesized that variability in IQ, gait speed, and SRT-hearing would be predicted by overlapping patterns of individual differences in the functional topography of these networks.

## MATERIALS AND METHODS

### Study Design and Participants

Participants were members of the Dunedin Study, a representative birth cohort (N = 1037; 91% of eligible births; 52% male) born between April 1972 and March 1973 in Dunedin, New Zealand (NZ) and eligible based on residence in the province and participation in the first assessment at age 3 years (Poulton et al., 2015; 2022). The cohort represented the full range of socioeconomic status in the general population of NZ’s South Island and, as adults, matches the NZ National Health and Nutrition Survey on key adult health indicators (e.g., body mass index, smoking, physical activity, physician visits) and the NZ Census of citizens the same age on educational attainment. The cohort is primarily white (93%). Assessments were carried out at birth and ages 3, 5, 7, 9, 11, 13, 15, 18, 21, 26, 32, 38, and most recently (completed April 2019) 45 years when 875 Study members completed neuroimaging. The NZ-HDEC (Health and Disability Ethics Committee) approved the Study and all Study members provided written informed consent. The concept and main analyses for this project were preregistered at rb.gy/34xv8a. All analyses were checked for reproducibility by an independent data analyst who used the manuscript, code, and an independent copy of the data to check all analyses.

### MRI Acquisition

Study members were scanned using a MAGNETOM Skyra 3T scanner (Siemens Healthcare GmbH) equipped with a 64-channel head/neck coil at the Pacific Radiology imaging center in Dunedin, New Zealand. High resolution T1-weighted images were obtained using an MP-RAGE sequence with the following parameters: TR = 2400 ms; TE = 1.98 ms; 208 sagittal slices; flip angle, 9°; FOV, 224 mm; matrix =256×256; slice thickness = 0.9 mm with no gap (voxel size 0.9×0.875×0.875 mm); and total scan time = 6 minutes and 52 seconds. Functional MRI (fMRI) data were collected using 72 interleaved axial T2-weighted functional slices with 3-fold multi-band accelerated echo planar imaging (TR=2000ms; TE=27ms; flip angle = 90 degrees; field of view=200mm; voxel size=2mm isotropic; slice thickness=2mm without gap). Resting-state and task fMRI data were collected as follows: (1) resting-state with eyes open and a gray screen displayed (8:16 min; 248 TRs), (2) emotional face processing task (6:40 min, 200 TRs), (3) color Stroop task (6:58 min, 209 TRs), (4) monetary incentive delay task (7:44 min, 232 TRs), and (5) episodic memory task (5:44min, 172 TRs). Details for each of the four tasks can be found in the **Supplemental Methods**. Twenty Study members completed the entire scanning protocol a second time (mean days between scans = 79 days) allowing for calculation of test-retest reliability of all our MRI-derived measures.

### fMRI Preprocessing

Resting-state and task fMRI data were concatenated into one time series to derive estimates of General Functional Connectivity (GFC), which we have shown enhances test-retest reliability and improves prediction of behavior in the Dunedin Study and other datasets (Elliott et al., 2019). The use of GFC was further motivated by strong convergent evidence that functional networks are largely invariant to task conditions and task-derived functional networks are similar to those derived from resting-state data (Fair et al., 2007; Gratton et al., 2018). Further analyses validating GFC in this dataset and others have been previously described (Elliott et al., 2019). Briefly, data were analyzed using the Human Connectome Processing minimal preprocessing pipeline (Glasser et al., 2013). T1-weighted anatomical images were skull-stripped, intensity-normalized, and nonlinearly warped into a study-specific average template in MNI space (Avants et al., 2008; Klein et al., 2009). Functional time-series data were despiked, slice-time-corrected, and realigned to the first volume in the time-series using AFNI (Cox, 1996). To limit distortion caused by in-scanner head motion, motion regressors were generated using 6 motion parameters and their first derivatives (to account for nonlinear effects) for a total of 12 motion regressors. Five components from white matter and cerebrospinal fluid were extracted (Behzadi et al., 2007) and used as nuisance regressors along with the mean global signal. Images were bandpass filtered to retain frequencies between 0.008 and 0.1 Hz. To reduce the influence of motion-related artifacts, we excluded all high-motion participants and adhered to strict scrubbing of motion-infected timepoints. We investigated a range of framewise-displacement cutoffs using QC-RSFC plots to determine the optimal threshold for removing motion artifacts as recommended by (Power et al., 2014). We selected 0.35mm framewise-displacement and 1.55 standardized DVARS as exclusion thresholds. Nuisance regression, bandpass filtering, censoring, and global-signal regression were performed using AFNI’s 3dTproject. For task data, functional connectivity due to signal evoked by task structure was removed using a Finite Impulse Response model (Fair et al., 2007). Finally, time series data were then projected to a two-dimensional fs_LR32k cortical surface space made up of ∼30,000 points or “vertices” that can be convoluted to improve anatomical correspondence between people (Van Essen et al., 2012).

Of the 875 Study Members who underwent MRI scanning, 769 were included in the current analyses after quality control procedures. 62 Study Members were excluded for excess head motion, 14 were excluded based on visual inspection of irregularities in a 360 ROI x 360 ROI functional connectivity matrix, 10 were missing one or more functional scan (i.e. rest and four tasks), 8 were excluded due to missing 3D-FLAIR sequences, 7 were excluded as MRI data were acquired with a 20-channel head coil instead of the 64-channel head coil to accommodate larger head circumferences, 4 were excluded due to incidental neurological findings, and 1 was excluded due to a missing fieldmap.

### Multi-session Hierarchical Bayesian Modeling (MS-HBM)

We implemented MS-HBM per the strategy of Cui et al. (2020), who applied this method in a dataset with a similar amount of fMRI data to the Dunedin Study. Broadly, MS-HBM defines several parameters that are modeled in a variational Bayes expectation maximization algorithm to estimate network labels across the cortical surface. Parameters required for this method are inter-region variability, inter-subject variability, and intra-subject variability as well as two tuning parameters which control the importance of the group average parcellation and smoothness.

To estimate inter-regional variability, MS-HBM first generates binarized connectivity profiles for 1483 equally spaced vertices across the cortical surface. Binarized profiles were defined here as the 10% of vertices across the cortical surface with the strongest functional connectivity to each of these 1483 vertices. A group average network parcellation was then derived using the average binarized connectivity profiles of all participants. To estimate intra-subject variability with only one scan per Study member, time series data were split into two halves of 17:41 min each, as recommended by Kong et al. (2019). Next, to estimate inter-subject variability, we used a resampling method as we did not have a validation dataset and computing across all Study members was highly computationally expensive. We randomly resampled 50 sets of 200 Study members from the total datasets available. We calculated inter-subject variability across each of these 50 sets and averaged these estimates. Finally, we selected the following tuning parameters that were optimized using data from the Human Connectome Project (Kong et al., 2019) due to similarities in our preprocessing procedures: smoothness prior: *c* = 40; group spatial prior: *α*= 200. The smoothness prior (*c*) controls the penalty assigned for assigning two adjacent vertices to different networks. The group spatial prior (*α*)controls the weight of the group spatial prior. In other words, a parcellation estimated with a high group spatial prior will be more similar to the group average.

Given the above parameter estimates, MS-HBM generates an individual-specific parcellation from each Study member’s GFC time series data using a variational Bayes expectation maximization algorithm (Kong et al., 2019). We chose to parcellate brain function into 17 networks. This number represents a previously determined optimal solution to capture the structure of correlations between cortical regions and allows for ready comparison with existing literature (Yeo et al., 2011). Specifically, these 17 networks were derived by applying a clustering algorithm to functional connectivity strength between evenly distributed vertices across the cortex (Yeo et al., 2011). To evaluate how “well” the resulting individualized parcellations captured variation in BOLD signal, we calculated functional homogeneities for all individualized parcellations and compared them to homogeneities from template network parcellation (Yeo et al., 2011). Specifically, functional homogeneity is the average BOLD timeseries correlation between all pairs of vertices assigned to the same network. Therefore, higher homogeneity means that, on average, vertices within the same network are more functionally connected.

To estimate test-retest reliability we generated parcellations using data from each of two separate scanning sessions in a subset of 20 Study members who were scanned twice. We first estimated test-retest reliability of individual network surface areas using a two-way mixed-effects intraclass correlation coefficient (ICC) with session modeled as a fixed effect and subject as a random effect (Shrout and Fleiss, 1979). We also calculated test re-test reliability of whole brain topographic organization by computing the Dice coefficient between timepoints as well as all pairs of unique Study members. The Dice coefficient is a metric to determine the overlap between two sets of categorical features (Birn et al., 2013; Destrieux et al., 2010; Kong et al., 2019, 2021). The Dice coefficient can be calculated using the following formula:

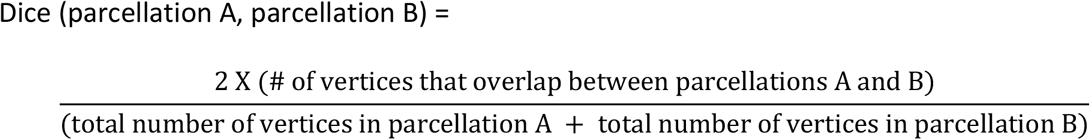

The Dice coefficient is equal to 1 if there is complete overlap between two parcellations and equal to 0 if there is no overlap between parcellations.

### Cognitive Function

Full-scale IQ was assessed at age 45 using the Wechsler Adult Intelligence Scale-IV (WAIS-IV; Wechsler, 2008). The WAIS-IV also yields indices of four specific cognitive domains: processing speed, perceptual reasoning, verbal comprehension, and working memory. Full-scale IQ was used in our primary analyses.

### Gait Speed

The gait speed of Study members was assessed using the GAITRite Electronic Walkway with 2m acceleration and deceleration before and after the walkway. Gait speed was assessed under three conditions: usual gait speed (walking at a normal pace from a standstill), maximum gait speed (walk as fast as safely as possible), and dual-task gait speed (walk at a normal pace while reciting alternate letters of the alphabet out loud). Study members completed each gait condition twice and the mean speed was taken for each condition. Gait speed was highly correlated across all three conditions (**Supplemental Results**). For our main analysis, we used the average of the usual gait speed and maximum gait speed conditions to remove any cognitive effects present in the dual-task condition. We also completed all gait-speed analyses using the mean of all three conditions to compare with previously published results from this cohort (Rasmussen et al., 2019). These results can be found in the **Supplemental Materials**.

### Listening Task

Study members completed the Listening in Spatialized Noise - Sentences test (LiSN-S) in a soundproof booth. Stimuli were presented using Sennheiser 215 headphones attached to a Mini PCM2704 external sound card. The LiSN-S generates a three-dimensional auditory environment in four different conditions. During the task, target sentences are superimposed with distractor sentences. The distractor sentences were presented at 55 decibels sound pressure level (dB SPL). Study members repeated target sentences out loud and were automatically scored according to the number of correct words in each sentence. The program began with target sentences presented at 62 dB SPL and intensity levels were adjusted according to performance: the intensity was adjusted down if > 50% of the words in a sentence were correct and adjusted up if <50% of the words were correct. The first several sentence presentations were considered practice, with each presentation lowered in 4 dB increments until performance dropped below 50% accuracy, after which increments decreased to 2 dB. Practice sessions were not included in the final scores.

The test condition continued until the average of the positive and negative-going reversals was ≥ 3 and the standard error of these midpoints was < 1 dB. If Study members did not reach this point, the test condition simply ended after 30 sentence presentations. Speech reception thresholds (SRT) were considered the lowest intensity a Study member could repeat 50% of words correctly. We used the low cue speech reception threshold, where the distractor speaker and target speaker had the same identity and were presented in the same location in the auditory environment, in our primary analyses. We selected low cue speech reception threshold as it is the most challenging condition and may have greater age-related variation. Crucially, lower scores on this measure indicate *better* hearing, as it reflects the intensity at which a person can successfully distinguish between distractor and target sentences. Further, we tested the other speech reception threshold from the LiSN-S task and measures of pure tone audiometry (see **Supplemental Materials**). We also tested results from an adaptation of the Digit Triplets Test (Van den Borre et al., 2021; King et al., 2011), a speech-in-noise task which uses white noise as a distractor instead of other speech (See **Supplemental Materials**). Results for these additional hearing measures can be found in the **Supplemental Results** and **Supplemental Figure S1**. We also repeated all analyses while controlling for overall hearing ability as measured by pure tone audiometry (**Supplemental Results, Supplemental Figure S2)**.

### Primary Analyses

We associated IQ, gait speed, and SRT-hearing with two broad features of functional topography: (i) total network surface area and (ii) spatial similarity. These are described in more detail below.

### Total Network Surface Area

Total network surface areas were calculated for all 17 functional networks by summing the number of vertices assigned to a given functional network in each Study member. As functional topography maps were generated after normalizing all subjects to fs_LR32k space, measures of network surface area already control for total cortical surface area. We conducted 17 univariate regressions between each functional network and each behavioral measure (IQ, gait speed, and SRT-hearing) while controlling for sex and in-scanner head motion (average framewise displacement). We used a Bonferroni corrected p value of .05/17 = 0.0029 to determine statistical significance.

To more stringently test associations observed with total network surface areas, we also trained linear ridge regression models using network surface areas and tested their ability to predict IQ, gait speed, and SRT-hearing in unseen data using a split-half scheme. Specifically, we used a 2-fold nested cross-validation scheme to train our model (Cui et al., 2020). The outer fold was used to estimate the generalizability of the model and the inner fold was used to optimize the penalty parameters. For each behavioral measure (IQ, gait speed, and SRT-hearing) we generated a rank ordering of Study members and placed odd-ranking Study members into the training set and even-ranking Study members into the test set (i.e., unseen data). This train/test set split was the outer fold. This approach to data splitting ensured that training and test sets would be closely matched on the behavioral outcome of interest.

We included an L2 regularization term to prevent model overfitting. This term was optimized during inner-loop cross-validation. Specifically, we again split the training set in half (training subset 1 and training subset 2). We then trained our ridge regression model using subset 1 to predict behavior in subset 2 using a variety of L2 regularization terms. We then repeated this procedure using subset 2 to predict subset 1. We calculated the accuracy of each prediction (Pearson’s *r*) and the mean absolute error (MAE) for each individual prediction. For each L2 term, we averaged the *r* and reciprocal of the MAE terms and summed these values to get an “accuracy” measure for each L2 term. The L2 term with the highest overall accuracy was carried forward to the outer fold step. Effects of sex and in-scanner head motion were regressed from our behavioral measures prior to model training. To prevent information leak, we regressed average FD and sex from the training set and applied the resulting beta weights to the test set. This approach ensures that our model only uses information from the training set, including covariate regression, when calculating predictions in the test set.

We employed a permutation method to test our predictions for statistical significance. Specifically, we shuffled the behavioral scores within the training set 1,000 times and recomputed the prediction strength each time. We considered observed predictions statistically significant if they had prediction accuracy above the 95th percentile of null predictions and prediction error below the 5th percentile of null predictions. Finally, to ensure prediction accuracies were not due to our data splitting procedure, we generated 100 random outer fold splits and generated predictions for each of these random splits to compare with our estimates derived from rank-order splitting.

We utilized the Haufe transform to assess for relative feature importance in our multivariate prediction model (Haufe et al., 2014). Because multivariate regression models involve transforming predictor variables from a high-dimensional feature space into a lower-dimensional feature space, the multivariate beta weights cannot be interpreted as relative feature importance. Instead, we calculated the covariance between each predictive feature and outcome variables. These covariances can be thought of as “relative feature importance” and can allow for interpretation of our predictive model. Prior to Haufe transformation, we standardized total surface area measures and regressed the effects of sex and in-scanner head motion from each of our behavioral variables. We then calculated covariance between each network’s standardized surface area and each predicted behavioral score. To maintain a consistent scale across behavioral measures, we divided each covariance measure by the variance of respective behavioral measures. To ensure that our feature importance estimates were not driven by idiosyncrasies in the data splitting procedure, we computed feature importance estimates across the aforementioned 100 random outer fold data splits. We report the average feature importance across these 100 predictions. Notably, we performed this analysis only using Study members in the training set to better interpret our trained regression model.

### Spatial Similarity

To test behavioral associations with the spatial layout of networks, we again used a split-half approach to train kernel ridge regression models using brain wide topographic similarity and tested its ability to predict behavior in held-out data. These models predict behavior of a held-out Study member as the weighted average of associations observed in Study members from the training set. The weights were determined by that held-out Study member’s brain-wide topographic similarity to each of the training Study members, as quantified by the Dice coefficient (Chen et al., 2022; Kong et al., 2019). We used the Dice similarity coefficient between Study members to estimate which Study members should show the most similarity in behavior. In other words, if two Study members have similar brain-wide functional topography, our models would predict that they have similar behavior. Additional details of this model can be found in the **Supplemental Methods**. We used the same 2-fold nested cross-validation approach as above to train and test the regression models using spatial similarity. We again used similar permutation methods to test our predictions for statistical significance and for robustness to data splitting procedures.

### Secondary Analyses – Regional Prediction

To assess for specific regional contributions to prediction accuracies, we assessed the accuracy of our kernel ridge regression models while training them using only topographic similarity from within 360 cortical parcels (Glasser et al., 2016). By using a parcellation that is not based on our functional connectivity maps, we are able to extract regional patterns of functional topography across the cortex. Due to the misalignment of these parcellations, each of the 360 parcels contain individual-level variation in functional network boundaries. Thus, regional predictions of behavior are based only on the topography within a specific region of the cortex. This procedure allowed us to test for qualitative similarity of the cortical regions most important for the predictions of IQ, gait speed, and SRT-hearing.

To formally test for correspondence between maps of parcel-wise prediction accuracies, we used a spatial permutation procedure known as a spin test (Váša et al., 2018). A spin test allows for tests of statistical significance by generating 10,000 random permutations of cortical surface data while preserving the spatial covariance structure of the data. A truly random shuffling of cortical data would create an unrealistically weak null distribution by generating biologically improbable distributions across the cortex (Alexander-Bloch et al., 2018). Thus, a spin test is a more conservative test of spatial correspondence. Correlations between two brain maps were considered statistically significant if they were higher than the 95th percentile of the set of null distributions generated through the spin test.

## RESULTS

### Cohort Characteristics

Attrition analyses revealed no significant differences in either childhood IQ or socioeconomic status between the full cohort, those still alive, those seen at age 45, or those scanned at age 45 (**Supplemental Methods, Supplemental Figure S3-S4**). Of the 875 Study members completing the neuroimaging protocol at age 45, 769 had GFC data passing quality control that were included in the primary analyses. Study members with usable GFC data did not significantly differ from the full age 45 sample in IQ at age 45 (*t* = 1.44, *p* = .15) or sex distribution (*X*^*2*^ = 0.21, *p* = 0.88). Thus, the current analyses continue to reflect effects in a population-representative cohort.

IQ, gait speed, and SRT-hearing were all normally distributed in the subsample of 769 Study members with high-quality GFC data (**Supplemental Figure S5A**). IQ, gait speed, and SRT-hearing were all significantly correlated (IQ and gait: r = .38, p < .001; IQ and SRT-hearing: r = -.31, p < .001; gait and SRT-hearing: r = -.20, p < .001; **Supplemental Figure S5B)**. This is consistent with previously reported associations between these variables in the full Dunedin Study cohort at age 45 (Rasmussen et al., 2019).

### Functional Topography Homogeneity and Test-Retest Reliability

Consistent with previous work (Cui et al., 2020; Kong et al., 2019, 2021), we found that individualized parcellations showed reliable differences between people and boosted average functional homogeneity (i.e., the average correlation in BOLD signal between all pairs of vertices within each network) compared to the template parcellation. Each individual Study member showed higher homogeneity from their individualized parcellation compared to the template parcellation (**Figure 1, Supplementary Figure S6**).

**Figure 1.**
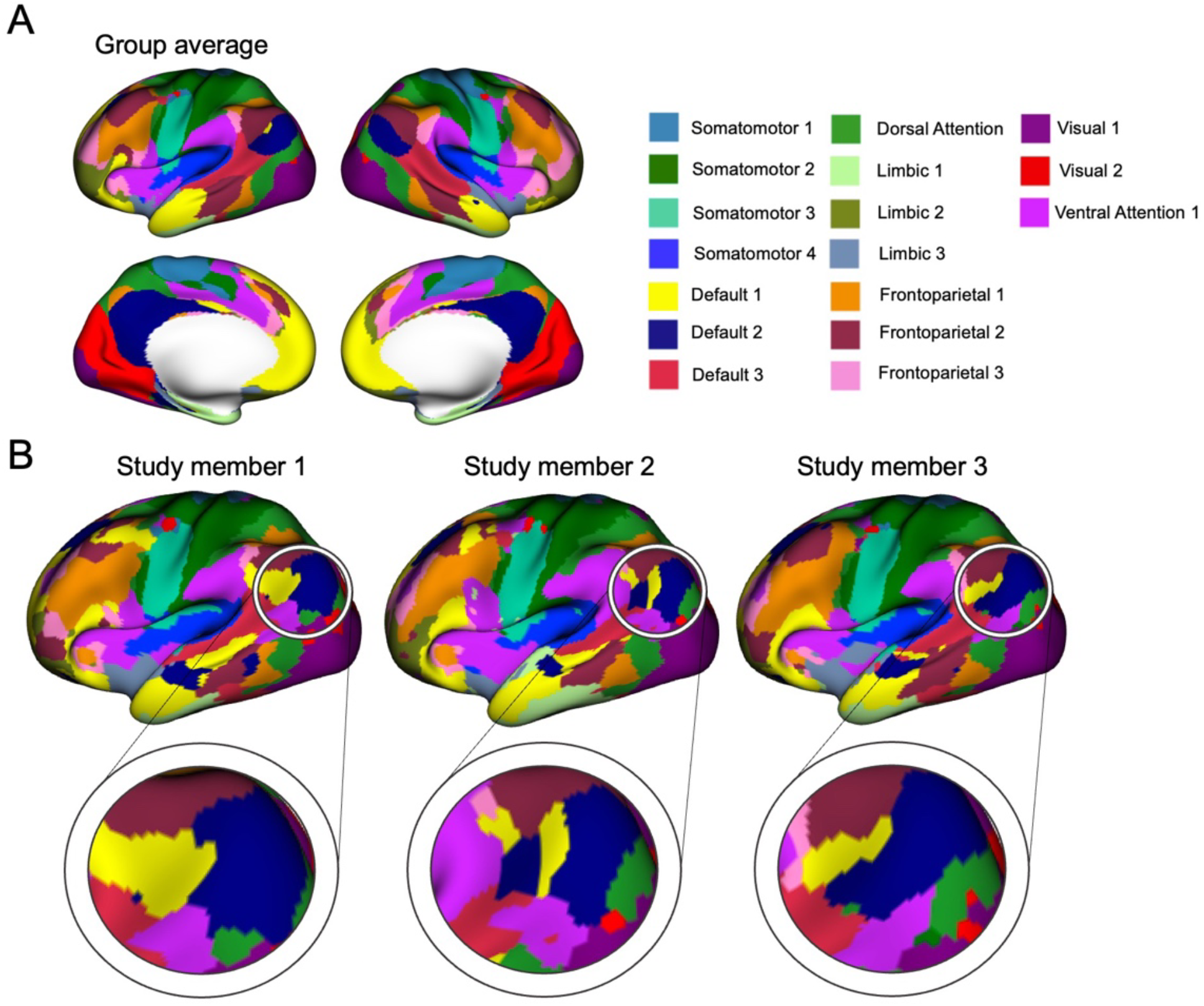
Example of functional topography in the Dunedin Study cohort. **A**. Group average functional parcellation. Legend shows colors corresponding to each of 17 neocortical networks. **B**. Example of functional topography variation in 3 Study members. Right hemisphere is shown to portray overall group correspondence while a region of the temporo-parietal junction is highlighted to show individual variation in topography.

High quality GFC data were available from 19 of the 20 Study members who repeated the scanning protocol. We found that total network surface areas showed fair test-retest reliability (mean network ICC = .47). We further observed strong within-person similarity between timepoints (Dice coefficient = 0.81), which was greater than between-person similarity (Dice coefficient = 0.72; **Supplemental Figure S7**). These results are consistent with previous test-retest analyses of functional topography in other datasets (Cui et al., 2020; Kong et al., 2019, 2021). Thus, our maps capture reliable individual-level features of functional topography.

### Total Network Surface Area

We first tested associations between total network surface area for each of the 17 functional networks and IQ, gait speed, and SRT-hearing (**Figure 2A**). IQ and gait speed but not SRT-hearing exhibited a broadly similar pattern of associations with network surface area. Notably, higher IQ and faster gait speed were associated with relatively larger default mode network 1, but relatively smaller default mode network 2 and limbic networks. No statistically significant associations with SRT-hearing were observed. A complete list of results is presented in **Supplemental Table S1**.

**Figure 2.**
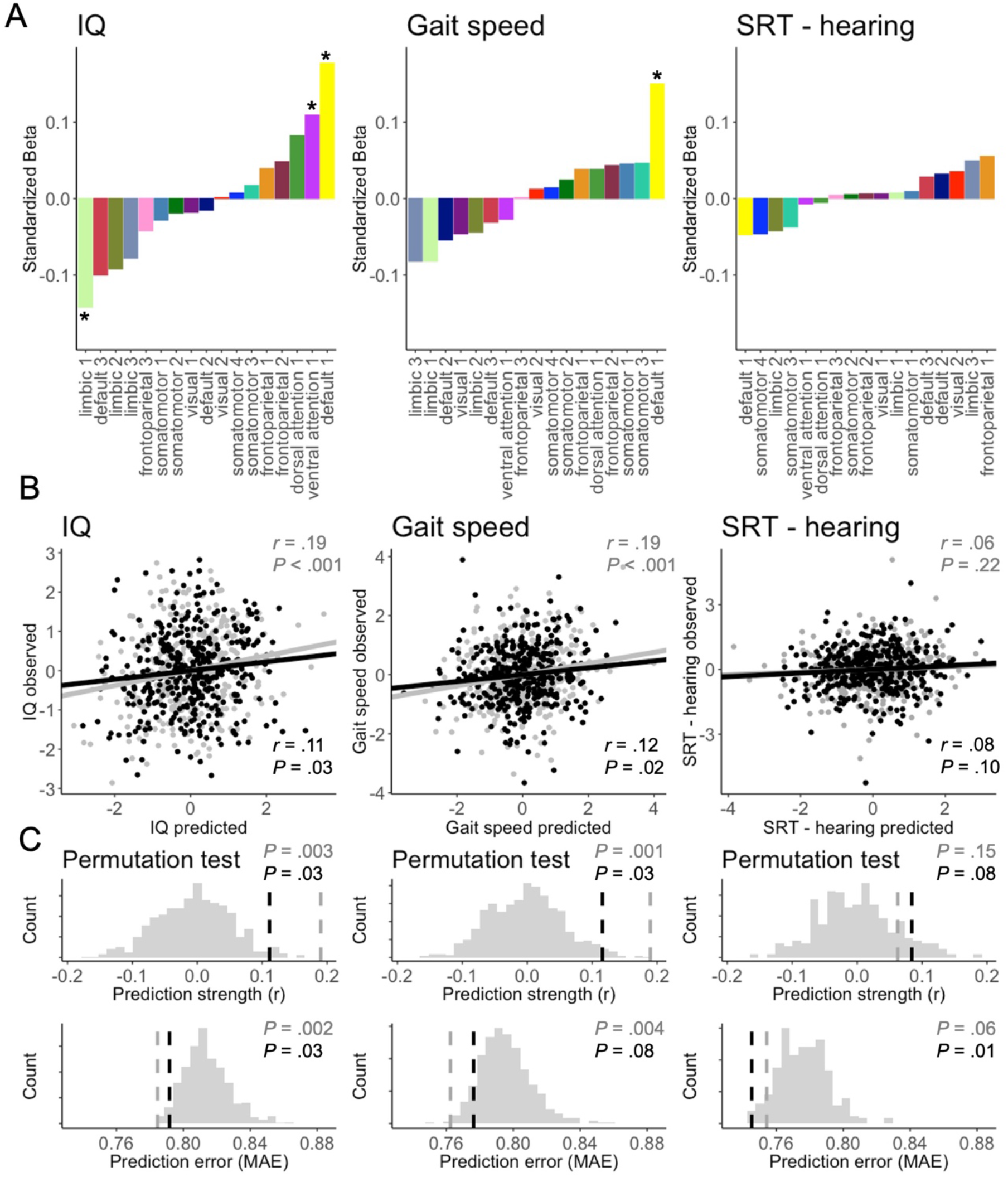
Network surface area analyses. **A**. Results from univariate correlations between network surface area and IQ, gait, and SRT-hearing. Colors correspond to functional networks identified in Figure 1A. Correlations marked with a star are statistically significant after Bonferroni correction across the 17 comparisons. **B**. Results from surface area prediction models for IQ, gait speed, and SRT-hearing. Scatterplots represent the correlation between observed scores and predicted scores according to ridge regression models. Fold 1 is shown in gray while fold 2 is shown in black. **C**. Permutation tests of observed predictions compared to 1,000 null predictions. (Top panel) Histograms show null distribution of prediction strengths and vertical lines show the observed prediction strengths. (Bottom panel) Histograms show the null distribution of prediction error and vertical lines show the observed prediction error. Fold 1 is shown in gray vertical lines while fold 2 is shown in black.

To gauge the similarity between profiles of associations, we computed correlations between resulting beta weights with IQ, gait speed, and SRT-hearing. While the pattern of associations appeared highly consistent between IQ and gait speed (r = .76, p = <.001), there was no such consistency between IQ and SRT-hearing (r = -.31, p = .22), or between gait speed and SRT-hearing (r = -.40, p = .11).

In complementary analyses, ridge regression models trained using total network surface areas (henceforth: surface area prediction models) were able to predict variation in IQ (fold 1: r = .19, p < .001; fold 2: r = .11, p = .03) and gait speed (fold 1: r = .19, p < .001, fold 2: r = .12, p = .02), but not SRT-hearing (fold 1: r = .06, p = .22, fold 2: r = .08, p = .10; **Figure 2B**). Our permutation procedure revealed that our predictions were significantly higher than would be expected by chance for IQ (fold 1: p_perm_ = .003, fold 2: p_perm_ = .03) and gait speed (fold 1: p_perm_ = .001, fold 2: p_perm_ = .03), but not for SRT-hearing (fold 1: p_perm_ = .15, fold 2: p_perm_ = .08). The predictions for IQ also had less error than would be expected by chance (fold 1: p_perm_ = .002, fold 2: p_perm_ = .03). Predictions for gait speed (fold 1: p_perm_ = .004, fold 2: p_perm_ = .08) and SRT-hearing also had relatively low error (fold 1: p_perm_ = .06, fold 2: p_perm_ = .01) (**Figure 2C**). These predictions were robust to variation in data-splitting procedures (**Supplemental Figure S9**).

Lastly, Haufe-transformed predictive feature scores derived for each functional network were broadly consistent between IQ and gait speed (r = .74, p = .001), but not between IQ and SRT-hearing (r = -.28, p = .26), or gait speed and SRT-hearing (r = -.37, p = .14). In addition, surface areas for default mode network 1 had strong positive feature importance for both IQ and gait, while limbic networks had strong negative feature importance for both IQ and gait (**Supplemental Figure S10**).

### Spatial Similarity

Next, we used a more fine-grained approach to testing the associations between functional topography and behavior. Using total network surface areas is a somewhat coarse way of testing associations with functional topography, as it does not consider the *shape* of functional networks. Furthermore, this coarse measure of total network surface area had only fair test re-test reliability in our dataset. Thus, the following analysis is based on the *shape* of individualized functional networks - a more detailed and reliable strategy of capturing functional topography. For this analysis, we again used a split-half approach to train kernel ridge regression models, this time using brain-wide topographic similarity as calculated using the Dice coefficient (henceforth: spatial similarity prediction models; **Figure 3A**). We found that these spatial similarity prediction models were able to predict IQ (fold 1: r = .34, p < .001, fold 2: r = .23, p < .001) and gait speed (r = .19, p < .001, fold 2: r = .10, p = .04), but not SRT-hearing (fold 1: r = .03, p = .58, fold 2: r = .05, p = .32) in held-out Study members (**Figure 3B**). Prediction was stronger than would be expected by chance for IQ (fold 1: p_perm_ < .001, fold 2: p_perm_ < .001) and gait speed (fold 1: p_perm_ < .001, fold 2: p_perm_ < .03), but not SRT-hearing (fold 1: p_perm_ = .28, fold 2: p_perm_ = .14). In addition, predictions for IQ (fold 1: p_perm_ < .001, fold 2: p_perm_ < .001) had less error than would be expected by chance. Gait speed had relatively low levels of error (fold 1: p_perm_ = .001, fold 2: p_perm_ = .23), but SRT-hearing prediction had error similar to the null distribution (fold 1: p_perm_ = .64, fold 2: p_perm_ = .09) (**Figure 3C**). Finally, these results were robust to variation in data splitting (**Supplemental Figure S11**). As has been observed previously (Kong et al., 2019, Cui et al., 2020), predictions using similarity tended to have stronger accuracy and less error compared to predictions using only total surface area (**Supplemental Figure S12**).

**Figure 3.**
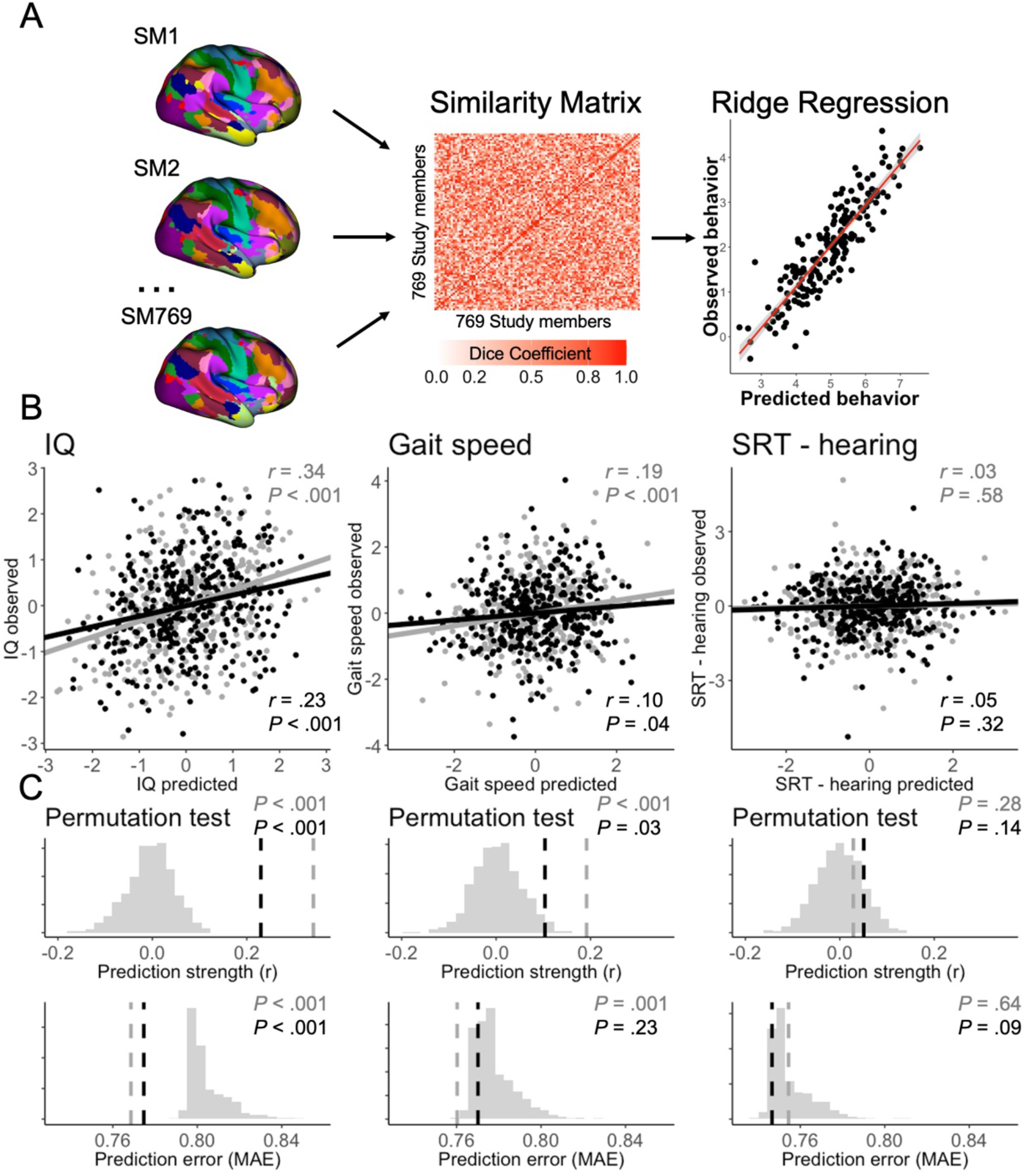
Whole brain spatial similarity analyses. **A**. Schematic depicting the organization of our ridge regression model. Brain-wide similarity (Dice coefficient) was calculated between all unique pairs of Study members and stored in a similarity matrix. This matrix was then used to train our ridge regression model to predict IQ, gait speed, and SRT in unseen Study members. **B**. Results of spatial similarity prediction models for IQ, gait speed, and SRT-hearing. Scatterplots show the correlations between observed scores for IQ, gait speed, and SRT-hearing and the predicted scores according to our ridge regression models. Fold 1 is shown in gray while fold 2 is shown in black. **C**. Permutation testing of spatial similarity prediction models. (Top panel) Histograms show the null distribution of prediction strengths from 1,000 null predictions. Vertical lines represent observed prediction strengths. (Bottom panel) Histograms show the null distribution of prediction error from 1,000 null predictions. Vertical lines represent observed prediction error. Fold 1 is shown in gray while fold 2 is shown in black

### Regional Prediction

To identify specific regional contributions to our observed predictions, we repeatedly assessed the accuracy of our spatial similarity prediction models while training them using only functional topography from within each of 360 anatomically derived cortical parcels (**Figure 4A**). Specifically, we calculated the Dice coefficient similarity between all pairs of participants for each cortical parcel. We then used each of these 360 similarity matrices to train our spatial similarity prediction models. Each resulting prediction strength thus indicates how well functional topography from within that anatomical parcel alone can predict behavior. This procedure also allowed us to test for qualitative similarity of the cortical regions most important for the predictions of IQ, gait speed, and SRT-hearing. Broadly, the strongest regional predictions for IQ and gait speed reflected variability in the functional topography of the temporoparietal junction and superior temporal gyrus, regions assigned to various default mode subnetworks in our group average, as well as lateral frontal cortex. SRT-hearing was modestly predicted from variability in lateral frontal and temporoparietal parcels (**Figure 4B**).

**Figure 4.**
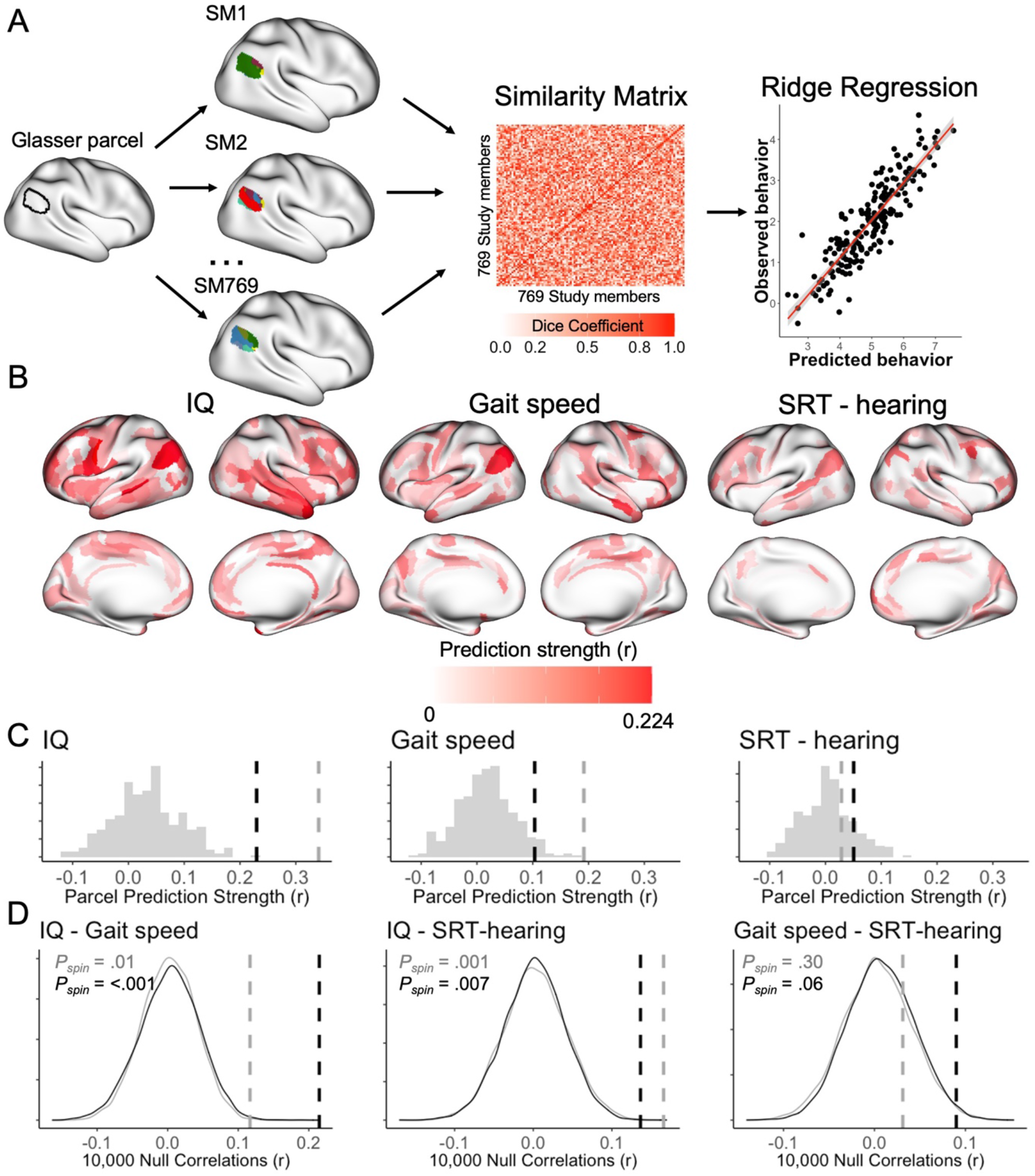
Regional prediction analyses. **A**. Schematic depicting our method for regional prediction. For each of the 360 Glasser parcels, we generated a new similarity matrix based on topographic similarity between all unique pairs of Study members within only that Glasser parcel. We then tested the ability for topography in that parcel alone to predict IQ, gait speed, and SRT-hearing. **B**. Prediction strengths for 360 Glasser parcels for IQ, gait speed, and SRT-hearing. Darker red indicates stronger prediction based on topography within that parcel. Parcels that had negative prediction values (i.e., no predictive strength) have no color. **C**. Comparison of regional predictions with whole brain predictions. Histogram shows distribution of predictions from each parcel and the vertical lines show prediction strength based on the whole brain. Fold 1 is shown in gray while fold 2 is shown in black. **D**. Cross-parcel correlations in predictions for IQ, gait speed, and SRT-hearing (vertical lines), overlaid on null density plots generated with spin tests. Fold 1 is shown in gray while fold 2 is shown in black. Leftmost plot shows the correlation between parcel-wise prediction of IQ and gait, middle plot shows the correlation between parcel-wise prediction of IQ and SRT-hearing, and rightmost plot shows the correlation between parcel-wise prediction of gait speed and SRT-hearing.

We also compared the prediction strength of each individual parcel with the prediction strength using global functional topography (i.e., across the entire cortex). We observed that the prediction of IQ based on global topography was stronger than prediction from the topography of any single parcel, while a small number of parcels predicted gait speed more strongly than did global topography. There was no difference between prediction strengths of regional or global topography for SRT-hearing (**Figure 4C**).

We used a spin test to formally evaluate correspondence between regional prediction maps for IQ, gait speed, and SRT-hearing. This test created 10,000 random permutations of values in the 360 cortical parcels while preserving their overall spatial covariance structure, thus generating a more realistic and stricter null distribution to test the observed association (Vasa et al., 2018). The spatial patterns of regional prediction for gait speed and SRT-hearing both modestly aligned with IQ (IQ-gait speed fold 1: r = .12, p_spin_ = .01; IQ-gait speed fold 2: r = .21, p_spin_ < .001; IQ-SRT-hearing fold 1: r = .17, p_spin_ < .001; IQ-SRT-hearing fold 2: r = .14, p_spin_ = .007). However, regional prediction patterns for gait speed and SRT-hearing did not align with each other (gait speed-SRT-hearing fold 1: r = .03, p_spin_ = .30; gait speed-SRT-hearing fold 2: r = .06, p_spin_ = .09; **Figure 4D**).

## DISCUSSION

We used a recently developed approach to reliably capture individual differences in the organization of functional networks across the neocortex (i.e., functional topography) in a large population-representative birth cohort now in midlife. We then leveraged this information to help better understand commonly observed correlations between seemingly disparate abilities, namely cognitive functioning as captured by IQ, and sensorimotor functioning as captured by gait speed and SRT-hearing. First, we found evidence for considerable variation in the topographical organization of common functional neocortical networks across this population-representative cohort, bolstering prior work with samples of convenience (Kong et al., 2019, Cui et al., 2020; Keller et al., 2022). Next, we found evidence for shared variation in the functional topography associated with IQ and gait speed but not SRT-hearing. While IQ and gait speed would appear to be quite different behaviors, they mapped onto highly similar aspects of functional topography. This suggests that covariation between cognitive and motor abilities during midlife is partially driven by shared variation in the functional integrity of widely-distributed and rather highly circumscribed neocortical networks.

More specifically, we found that higher IQ and faster gait speed both mapped onto increased surface area of higher-order functional cortical networks typically associated with cognitive functions and not lower-order somatomotor cortical networks typically associated with motor functions. This is in line with prior findings that gait speed in older adults most strongly correlates with the functional connectivity strength of higher-order functional networks (Lo et al., 2017; Yuan et al., 2015). However, these prior studies utilized functional *connectivity* rather than functional *topography* so we cannot directly compare our results to these prior studies. In our analyses, variation in IQ and gait speed most strongly reflected the functional topography of the default mode network. Thus, slower gait speed in midlife appears to reflect changes in default mode network organization. Notably, the default mode network is thought to be critical during aging, cognitive decline, and dementia onset (Buckner et al., 2005). Specifically weaker functional connectivity of the default mode network have been associated with aging, cognitive decline, and poorer health in adults (Sambataro et al., 2010; Smith et al., 2015; Staffaroni et al., 2018). The observed topographic patterns of associations with IQ and gait speed may index poorer overall health at midlife that may predispose people to accelerated biological aging (Elliott et al., 2021), and could reflect the early signs of aging itself. This would align with the ‘last in, first out’ hypothesis of brain aging (Douaud et al., 2014), which posits that the last brain networks to develop during early life (i.e., default mode network) are the first to degrade during later life. Given the relative youth of our cohort compared to most aging studies (i.e., age 45 vs. age 65+), the cross-sectional associations with the default mode network could reflect early stages of aging that first manifest in higher-order, later-developing networks.

Notably, while we observed the strongest associations with the functional topography of the default mode network, our prediction models performed best when using information from all 17 neocortical networks. Thus, while the default mode network may have greater relative importance, the shared variance between IQ and gait speed reflects the functional organization of the entire neocortex.

In contrast, we did not observe similar associations between SRT-hearing and functional topography. This is somewhat surprising given that SRT-hearing was correlated with cognitive ability at a similar magnitude as gait speed in our cohort, and all three measures were similarly normally distributed. This null finding was true for SRT-hearing as well as other hearing variables and was robust to variation in overall hearing ability (see **Supplemental Materials** for additional analyses). It is important to note that very few Study members demonstrated clinically significant hearing loss at midlife. Age and peripheral hearing loss significantly contribute to SRT-hearing (Besser et al. 2015) and normal hearing has been shown to nullify the association between SRT-hearing and cognitive ability (Glyde et al., 2013). While many studies have found associations between sensory functioning and cognitive decline, these studies have typically focused on participants older than 65 years, who would typically have mild or greater hearing loss (Lin et al., 2011; Loughrey et al., 2018; Lindenberger & Baltes, 1994). A possible explanation for the absence of associations between SRT-hearing and functional topography is that sensory functioning is less closely tied to the organization of neocortical networks in midlife compared to cognitive and motor functioning. Indeed, when younger (25-69 years) and older (70-103 years) adults are directly compared, older adults show stronger associations between sensory and cognitive ability compared to younger adults (Baltes & Lindenberger (1997). Finally, prior research in the Dunedin Study has found that biological aging in midlife is more weakly associated with sensory functioning compared to cognitive and motor functioning (Elliott et al., 2021). Thus, if the profile of associations between functional topography, IQ, and gait speed does reflect midlife stages of biological aging, this profile may not yet reflect variation in hearing ability.

In addition to these novel findings, our analyses extend prior studies of functional topography in several ways. First, our individualized maps of functional topography had good test-retest reliability comparable with prior work (Kong et al., 2019, 2021), including with parcellation methods other than MS-HBM (Cui et al., 2020). Second, we found associations between functional topography and cognitive ability that have effect sizes and prediction performances that are similar to prior work in other datasets (Cui et al., 2020; Kong et al., 2019; Keller et al., 2022). Third, we replicated findings that measures of network spatial organization tend to outperform summary metrics of total network surface areas in predicting behavior (Kong et al., 2019). Taken together, these findings provide further evidence for the utility of functional topography as a novel technique for reliably capturing individual differences in brain function and their mappings out to behavior.

Our study is not without limitations. First, neuroimaging, gait speed assessment, and hearing measurements were only conducted at one time point, precluding longitudinal analyses. While we have previously found that gait speed is associated with longitudinal biological aging (Rasmussen et al., 2019), we are not yet able to describe longitudinal changes in gait speed itself, hearing ability, or functional topography. However, repeat testing of neuroimaging, gait speed, IQ, and hearing in this cohort will begin at age 52 in 2023. Second, the Dunedin Study cohort was established 5 decades ago, bringing an inherent limit on sample size. Thus, we faced power constraints typical of cross-sectional analyses of brain-behavior associations (Gratton et al., 2022; Marek et al., 2022). It is partially for this reason that we selected a strategy (MS-HBM) designed to reduce individual error inherent in traditional analyses of brain function. While our replications of prior brain-behavior findings from other large datasets are reassuring (Kong et al., 2019), we must await additional independent replications of our novel findings. Moreover, our population-representative dataset may generalize more readily than convenience samples that suffer from healthy volunteer bias. Finally, our preregistration and reproducibility-check strategies prevent p-hacking, which is known to contribute to reproducibility failures.

Taken together, our results present a profile of brain-wide functional organization that correlates with both cognitive and motor functioning during midlife. These convergent patterns across the functional topography of IQ and gait speed provide a plausible biological basis for why gait speed captures individual differences in midlife cognitive function (Rasmussen et al., 2019), overall health (Smith et al., 2015), and, potentially, early signs of accelerated aging in midlife. This suggests that gait speed may not be simply a measure of physical function but rather an integrative index of nervous system health.

## ACKNOWLEDGEMENTS AND DISCLOSURES

This research received support from US-National Institute on Aging grants R01AG069939, R01AG032282, and R01AG049789 and UK Medical Research Council grant MR/P005918/1. We thank the Dunedin Study members, Unit research staff, and Study founder Phil Silva. The Dunedin Multidisciplinary Health and Development Research Unit is supported by the New Zealand Health Research Council (Programme Grant 16-604) and New Zealand Ministry of Business, Innovation and Employment (MBIE). The Dunedin Unit is within the Ngāi Tahu tribal area who we acknowledge as first peoples, tangata whenua (translation: people of this land).

The authors report no biomedical financial interests or potential conflicts of interest.

## DATA AND CODE AVAILABILITY

Dunedin Study data are available via managed access (https://moffittcaspi.trinity.duke.edu/research). All code used in this analysis are available upon request.

