## Supplemental Materials for "Functional Topography of the Neocortex Predicts Covariation in Complex Cognitive and Basic Motor Abilities"

Ethan T. Whitman, et al.

#### SUPPLEMENTAL METHODS

##### Phase 45 Attrition Analysis

We conducted an attrition analysis using childhood intelligence quotient (IQ; **Figure S4**) and socioeconomic status (SES; **Figure S5**) to determine whether participants in the Phase 45 data collection were representative of the original cohort.

##### fMRI task details

*Emotion processing task.*

The task consists of four blocks of a perceptual face-matching task interleaved with five blocks of a sensorimotor control task. The Dunedin Study version of this task consists of one block each of fearful, angry, surprised, and neutral facial expressions presented in a pseudorandom order across participants. During face-matching blocks, participants view a trio of faces and select one of two faces (on the bottom) identical to a target face (on the top). Each face processing block consists of six images, balanced for gender, all of which were derived from a standard set of pictures of facial affect. During the sensorimotor control blocks, participants view a trio of simple geometric shapes (circles and vertical and horizontal ellipses) and select one of two shapes (bottom) that are identical to a target shape (top). Each sensorimotor control block consists of six different shape trios. All blocks are preceded by a brief instruction ("Match Faces" or "Match Shapes") that lasts 2 s. In the task blocks, each of the six face trios is presented for 4 s with a variable interstimulus interval (ISI) of 2-6 s (mean = 4 s) for a total block length of 48 s. A variable ISI is used to minimize expectancy effects and resulting habituation and maximize amygdala reactivity throughout the paradigm. In the control blocks, each of the six shape trios is presented for 4 s with a fixed ISI of 2 s for a total block length of 36 s.

*Stroop task.*

In this version of the Stroop task, participants identify the color of a target word in the center of a screen by selecting 1 of 4 identifier words. Selections are made by pressing 1 of 4 buttons on a button box, with each button matching an identifier word on the screen (e.g., index finger button 1 = identifier word on the far left, etc.). In congruent trials, targets were in colors congruent with the target words; in incongruent trials, targets are in colors incongruent with the targets. The four identifier words are in white in both conditions. After a 2 s fixation lead in, participants completed three 60 s blocks of congruent trials interleaved with three 60 s blocks of incongruent trials. Both conditions are followed by an 8 s fixation period for a total task time of 6 min and 50 s. Each block contains 12 trials, each consisting of 2 s stimuli, 1 s feedback, and a variable inter stimulus interval averaging 2 s.

*Monetary incentive delay (MID) task.*

This version of an event-related MID task consists of 12 eight second trials for each of 3 conditions, presented in a pseudo-random order, for a total of 36 trials. Conditions consist of potential $1 reward, potential $5 reward, and no monetary outcome. On reward trials, participants could win money by pressing a button during the presentation of a target. During each trial, participants see either a green dollar amount cue (reward conditions) or a white “$0” (neutral condition) (cue, 2 seconds), then fixate on an “x” as they wait for a variable interval (delay, 2250–3000 ms), and then respond with a button press to a white target triangle that appears for a variable length of time (target, 70-680 ms) with a button press. Feedback (feedback, 2 seconds), which follows the disappearance of the target, notifies participants of whether they had successfully responded to the target with either the word “HIT” or “MISS”, and indicates their cumulative total at that point. Initial task difficulty (i.e. duration of the target) is based on reaction times collected during a practice session before scanning, and an adaptive algorithm was employed to adjust the target’s duration such that each participant should succeed on ~66% of his or her target responses. Trials are separated by a variable (2-6 seconds) inter-trial interval. fMRI volume acquisitions are time-locked to the offset of each cue and thus were acquired during anticipatory delay periods.

*Memory encoding task.*

This task consists of the encoding and subsequent recall of novel face-name pairs. A distractor task (odd/even number identification) is interleaved between encoding and recall blocks to prevent maintenance of information in working memory. During each of four encoding blocks, subjects view six novel face-name pairs for 3.5 seconds each. During each of four recall blocks, subjects view six faces each presented for 2 seconds and immediately followed by an incomplete name fragment for 1.5 seconds during which they are required by forced-choice to determine if the fragment is correct or incorrect. A 1.5 second inter-trial interval is used during recall blocks. During each of four distractor blocks, subjects view six different numbers for 3.5 seconds each and are required to determine if the numbers are odd or even.

**Additional LiSN-S measures**

In addition to testing functional topographic associations with low-cue speech reception threshold (SRT), we also tested for associations with additional scores from the LiSN-S task. Briefly, the LiSN-S generates a three-dimensional auditory environment in four different conditions. During the task, target sentences are superimposed with distractor sentences. Study members repeated target sentences out loud and were scored according to the number of correct words in each sentence. Intensity levels were adjusted according to performance: the intensity was adjusted down if > 50% of the words in a sentence were correct and adjusted up if <50% of the words were correct. The test condition continued until the average of the positive and negative-going reversals was ≥ 3 and the standard error of these midpoints was < 1 dB. Speech reception thresholds (SRT) were considered the lowest intensity a Study member could repeat 50% of words correctly. We primarily used low-cue SRT in our analyses. Low-cue SRT represents the most challenging version of the task (both target and distractor sentences are from the same individual, both target and distractors sentences are in the same location). We also tested high-cue SRT, which represents SRT during the easiest listening condition (target and distractor sentences are spoken by different individuals, target and distractor sentences are presented at 90 degrees azimuth. In addition, we tested spatial advantage and talker advantage scores. Spatial advantage measures the difference in score gained when the target and distractor sentences are presented in different locations. Talker advantage measures the difference in score gained when the target and distractor sentences are spoken by different individuals. Finally, we tested total advantage score, which combines spatial and talker advantage scores.

**Pure tone audiometry**

To test our listening analyses for robustness to variation in hearing ability, we conducted all listening analyses again while controlling for hearing ability measured by pure tone audiometry. Pure tone audiometry was measured in a sound-attenuating booth (350 Series MaxiAudiology Booth by IAC Acoustics) and was administered using the Interacoustics Callisto Suite configured to the Interacoustics OtoAccess database. The stimuli were presented on an HP Envy laptop with Sennheiser HAD 300 headphones. Pure-tone stimuli of various frequency were presented in the following order: 1000 Hz, 2000 Hz, 4000 Hz, 8000 Hz, 12500 Hz, and 500 Hz. Intensity began at 40 decibels at hearing level for Study members with normal hearing. Intensity began at 60 decibels at hearing level for Study members using hearing aids. Participants responded when they heard a pure tone, according to the Huchson-Westlake procedure. Auditory thresholds were defined as the lowest intensity level that the individual responded to for two out of three presentations. Specifically, this threshold was determined using a standard down-10-up-5 technique. A mid-frequency pure tone average was calculated by averaging 1000, 2000, and 4000 Hz; and a high-frequency pure-tone average was calculated by averaging 8000 Hz and 12500 Hz. The results from “best ear” are reported.

**Digit Triplet Test**

We also tested scores derived from the New Zealand Hearing Screening Test (King, 2011; henceforth: Digit Triplet Test). This test consists of digits spoken in triplets with white noise in the background. Stimuli were delivered diotically using Sennheiser HAD 280 headphones. The high-definition audio volume was set at 70dB SPL (+/- 0.4 dB SPL). Study members listened to digits and reported what they heard to the examiner. Digits 1-9 were present, except for 7 as it is a di-syllabic word. The background noise was fixed at 65 dB SPL. Intensity of the spoken digits were adjusted according to the Study member's accuracy. If a Study member answered correctly, the intensity was decreased by 2 dB. If they answered incorrectly, the intensity was increased by 2 dB. After 27 triplets, the score was derived by taking the average of the previous 20 intensity levels.

**Spatial similarity prediction model**

We wish these models to predict behavior according to the complex spatial arrangement of functional networks, so our spatial similarity prediction models are not trained on any quantification of functional networks but rather on spatial similarity between people. Thus, this model implementation differs from the surface area prediction models and is described below. We implemented this model following the steps laid out in Kong et al. (2019).

Suppose we are trying to predict the behavioral measure for the *i*-th Study member ($y_i$) using their individualized parcellation ($l_i$). Assuming we have a set of behavioral measures for *N* Study members in the training set $\{y_1, y_2, \dots, y_N\}$ and individual parcellations for these same Study members $\{l_1, l_2, \dots, l_N\}$, we can predict $y_i$ using the following formula:

$$y_i = \sum_{j=1}^{N} \alpha_j K(l_{i,} l_j)$$

Where $K(l_i, l_j)$ is equal to the whole brain Dice similarity between the *i*-th and *j*-th Study members. In order to define $\alpha$ in the above equation, we can first define the set of behavior variables as $\boldsymbol{y} = [y_1, y_2, \dots, y_N]^T$ and the set of $\alpha$ values as $\boldsymbol{\alpha} = [\alpha_1, \alpha_2, \dots, \alpha_N]^T$. In addition, we can define **K** to be a symmetric $N$ x $N$ matrix with the Dice similarity for all pairs of Study members in the training set. As demonstrated by Kong et al., (2019), $\boldsymbol{\alpha}$ can be expressed as:

$$\boldsymbol{\alpha} = \boldsymbol{K}^{-1}\boldsymbol{y}$$

Therefore, we can predict the behavioral score of the $t$-th Study member in the test set using the following formula:

$$y_t = \boldsymbol{S}_t\boldsymbol{\alpha} = \boldsymbol{S}_t\boldsymbol{K}^{-1}\boldsymbol{y}$$

Where $\boldsymbol{S}_t$ is a vector of the Dice similarity coefficient between the $t$-th Study member in the test set and all Study members in the training set.

Conceptually, the predicted behavioral score can be thought of as a weighted average of the behavioral scores of the training set. The weights are determined according to the Dice similarity between the test subject and each of the training subjects. If the test subject has highly similar functional topography to a certain training subject, that training subject's behavioral score will be more heavily weighted. On the other hand, if the test subject has highly dissimilar functional topography to a certain training subject, that training subject's behavioral score will be weighted lower.

To reduce overfitting, we added an L2 regularization term with a tuning parameter, $\lambda$. Following Kong et al., (2019), this results in a new formula for $\boldsymbol{\alpha}$:

$$\boldsymbol{\alpha} = (\boldsymbol{K} + \lambda\boldsymbol{I})^{-1}\boldsymbol{y}$$

which ultimately produces our final formula for predicting behavior in subject $t$:

$$y_t = \boldsymbol{S}_t \boldsymbol{\alpha} = \boldsymbol{S}_t (\boldsymbol{K} + \lambda \boldsymbol{I})^{-1} \boldsymbol{y}$$

where ***I*** is an *N* x *N* identity matrix.

### SUPPLEMENTAL RESULTS

#### Gait conditions correlation

To validate our method of averaging gait speed across gait conditions (usual gait speed and maximum gait speed), we tested the correlations between these conditions in the subsample with useable fMRI data. As has been previously published (Rasmussen et al., 2019), these conditions were highly correlated with one another ($r = .45$, $p < .001$). In addition, both conditions were highly correlated with the dual-task gait condition ($r = .74$, $p < .001$; maximum gait and dual-task gait : $r = .44$, $p < .001$). While we used only usual gait speed and maximum gait speed to generate our composite gait score, our group has previously used an average of all three conditions (Rasmussen et al., 2019). The correlation between the two-condition composite used here and the three-condition composite used previously is extremely strong (r = .95, p < .001). Use of both composites yield highly similar correlations with functional topography (**Figure S8).**

#### Additional hearing analyses

We conducted further analysis of Study members' hearing abilities based on a variety of hearing measures described above. We did not observe any significant associations between any LiSN-S score, the Digit Triplet score, or pure tone audiometry score and any network surface area (**Figure S1A)**. We also did not observe significant prediction of any LiSN-S score, Digit Triplet score, or pure tone audiometry score surface area prediction models (**Figure S1B**), or when using our spatial similarity prediction models (**Figure S1C**).

In addition, we conducted these additional analyses while controlling for mid-frequency PTA to account for variation in peripheral hearing ability. Mid-frequency was selected as the control variable to approximate the frequency of speech. When controlling for mid-frequency PTA, we did not observe any significant associations between any LiSN-S score or the Digit Triplet score and any network surface area (**Figure S2A**). We also did not observe significant surface area prediction of any LiSN-S score or the Digit Triplet score while controlling for mid-frequency PTA (**Figure S2B**). Finally, we did not observe significant spatial similarity prediction for any LiSN-S score or the Digit Triplet score while controlling for mid-frequency PTA (**Figure S2C**).

**Table S1. Univariate analysis test statistics for IQ, gait speed, and SRT-hearing**

| | IQ | | Gait | | SRT - hearing | |
|---|---|---|---|---|---|---|
| **Functional Network** | **beta** | **p** | **beta** | **p** | **beta** | **p** |
| Default 1 | 0.18 | < .001 | 0.15 | < .001 | -0.05 | 0.198 |
| Default 2 | -0.02 | 0.661 | -0.05 | 0.112 | 0.03 | 0.351 |
| Default 3 | -0.10 | 0.005 | -0.03 | 0.394 | 0.03 | 0.432 |
| Dorsal Attention 1 | 0.08 | 0.022 | 0.04 | 0.296 | -0.01 | 0.899 |
| Frontoparietal 1 | 0.04 | 0.283 | 0.04 | 0.292 | 0.06 | 0.128 |
| Frontoparietal 2 | 0.05 | 0.179 | 0.04 | 0.225 | 0.01 | 0.863 |
| Frontoparietal 3 | -0.04 | 0.232 | 0.00 | 0.973 | 0.00 | 0.904 |
| Limbic 1 | -0.14 | < .001 | -0.08 | 0.022 | 0.01 | 0.846 |
| Limbic 2 | -0.09 | 0.008 | -0.04 | 0.205 | -0.04 | 0.230 |
| Limbic 3 | -0.08 | 0.031 | -0.08 | 0.024 | 0.05 | 0.175 |
| Somatomotor 1 | -0.03 | 0.427 | 0.05 | 0.208 | 0.01 | 0.801 |
| Somatomotor 2 | -0.02 | 0.594 | 0.02 | 0.506 | 0.01 | 0.888 |
| Somatomotor 3 | 0.02 | 0.643 | 0.05 | 0.206 | -0.04 | 0.298 |
| Somatomotor 4 | 0.01 | 0.837 | 0.01 | 0.709 | -0.05 | 0.203 |
| Ventral Attention 1 | 0.11 | 0.001 | -0.03 | 0.436 | -0.01 | 0.833 |
| Visual 1 | -0.02 | 0.619 | -0.05 | 0.210 | 0.01 | 0.866 |
| Visual 2 | 0.00 | 0.980 | 0.01 | 0.741 | 0.04 | 0.322 |

*Statistical significance was assessed using a Bonferroni correction across all 17 functional networks (.05 / 17 = .0029)

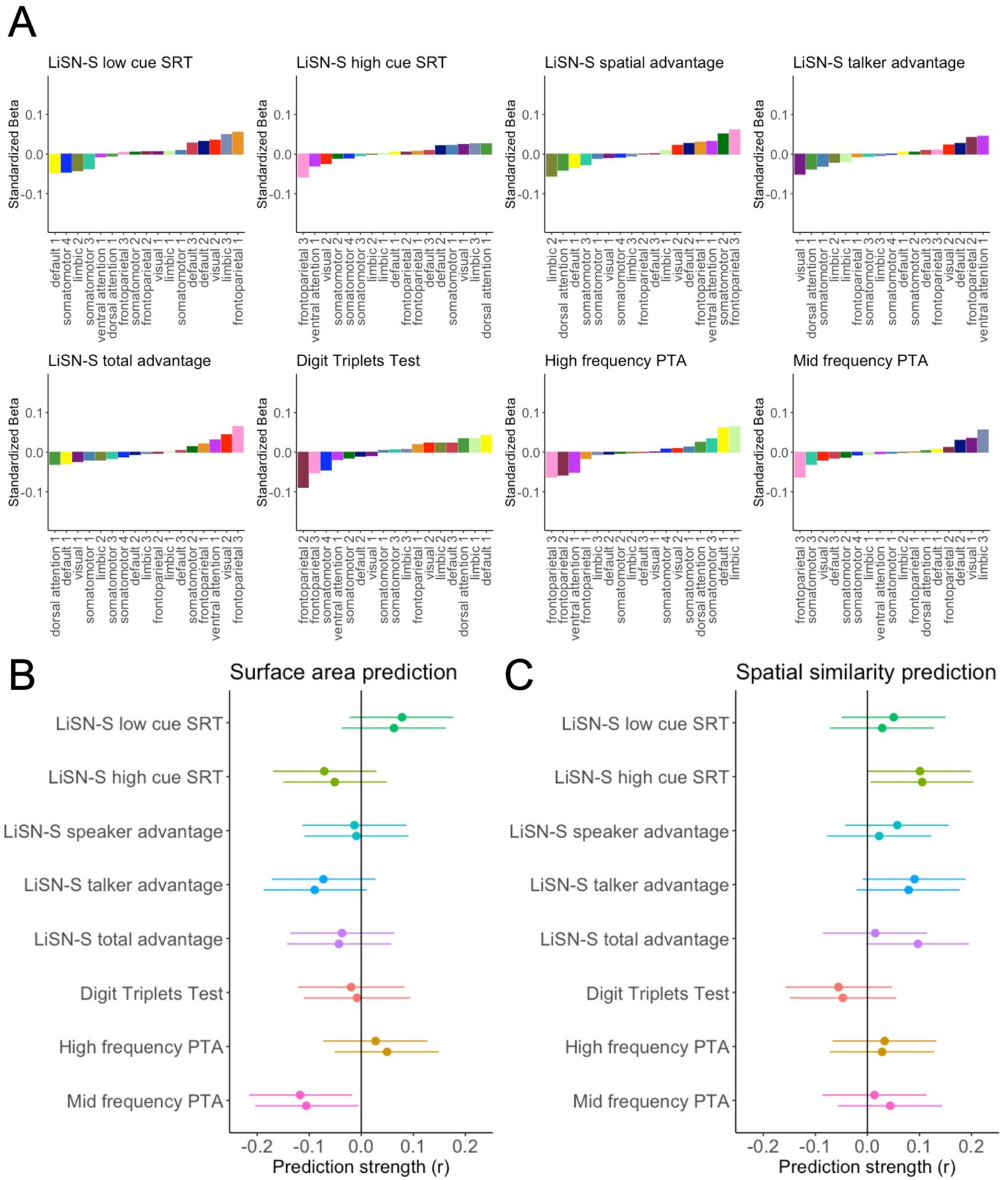

**Figure S1. Testing associations between functional topography and additional hearing variables.** Functional topography does not show associations with any scores from the LiSN-S, Digit Triplet Test score, or high- or mid-frequency PTA. A. Univariate associations between total network surface areas and additional hearing measures. No association was statistically significant after Bonferroni correction across 17 functional networks. B. Surface area prediction strengths for additional hearing measures. Both folds are shown. Error bars represent 95% confidence intervals. C. Spatial similarity prediction strengths for additional hearing measures. Both folds are shown. Error bars represent 95% confidence intervals.

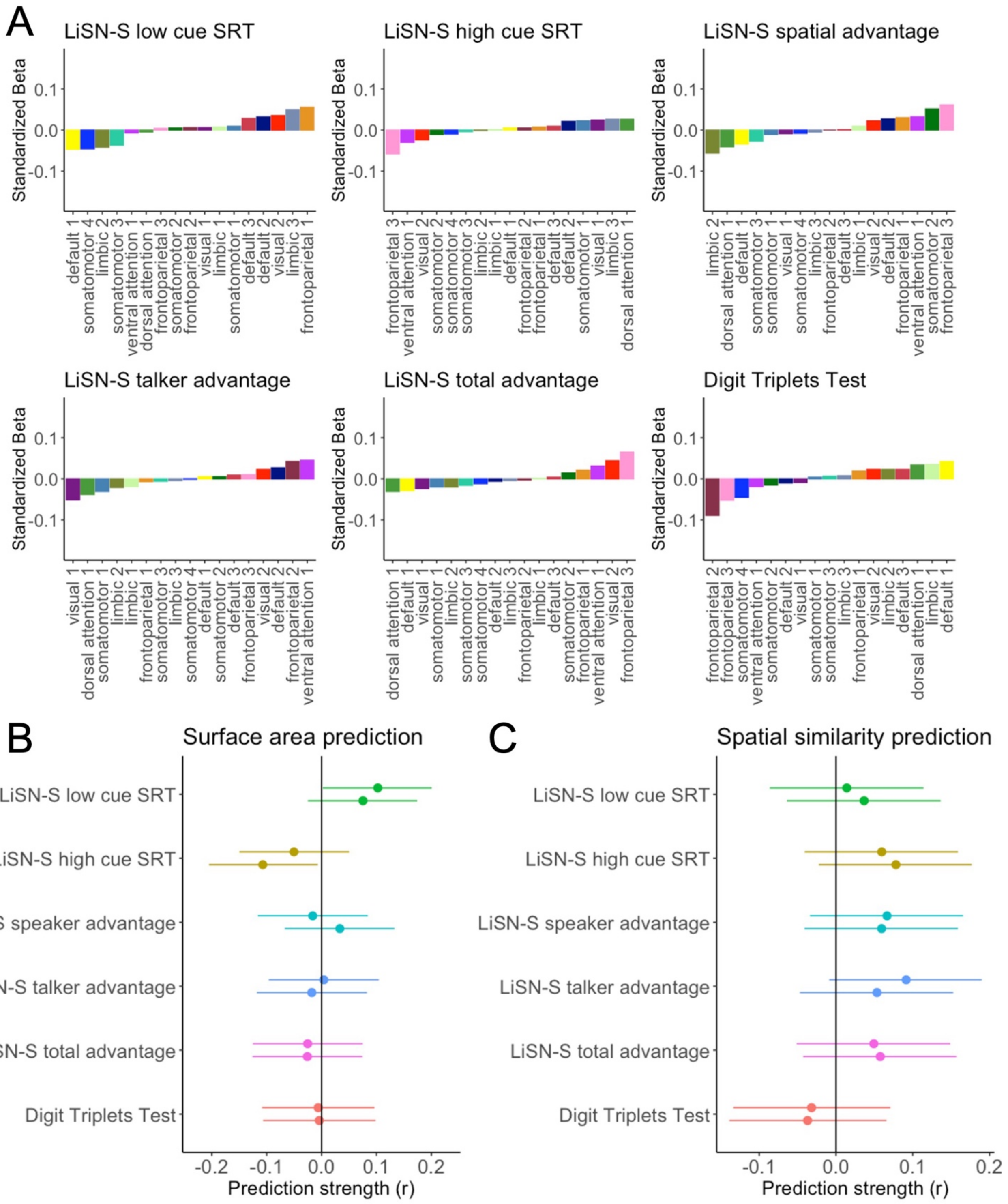

**Figure S2. Testing associations between functional topography and additional hearing variables while controlling for mid-frequency PTA.** Functional topography does not show associations with any scores from the LiSN-S or Digit Triplet Test score while controlling for mid-frequency PTA. A. Univariate associations between total network surface areas and additional hearing measures. No association was statistically significant after Bonferroni correction across 17 functional networks. B. Surface area prediction strengths for additional hearing measures. Both folds are shown. Error bars represent 95% confidence intervals. C. Spatial similarity prediction strengths for additional hearing measures. Both folds are shown. Error bars represent 95% confidence intervals.

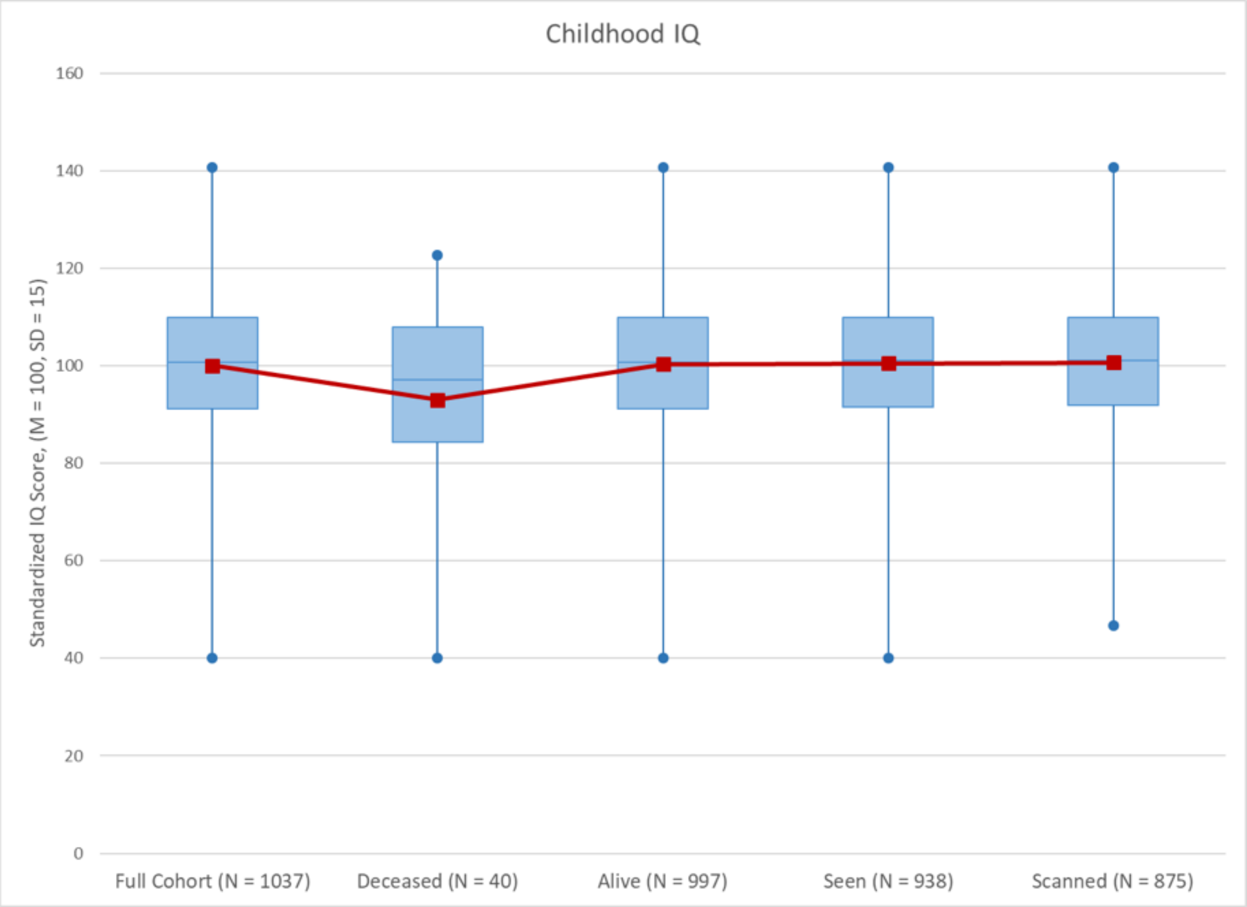

**Figure S3. Attrition analysis using IQ.** No significant differences in childhood IQ were found between the full cohort, those still alive, those seen at Phase 45 or those scanned at Phase 45. Those who were deceased by the Phase 45 data collection had significantly lower childhood IQ's than those who were still alive ($t = 2.09$, $p = 0.04$).

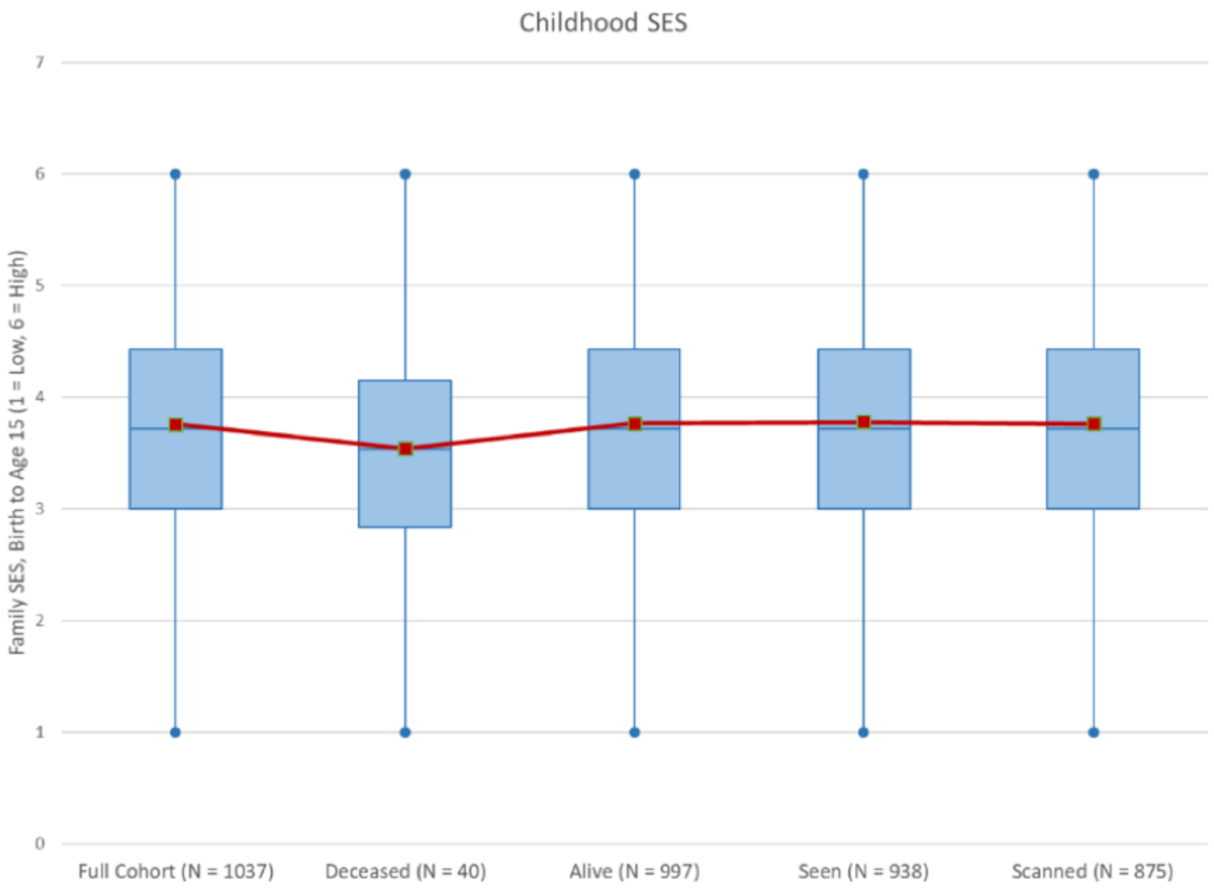

**Figure S4. Attrition analysis using SES.** No significant differences were found between the full cohort, those deceased, those alive, those seen at Phase 45 or those scanned at Phase 45 on childhood SES

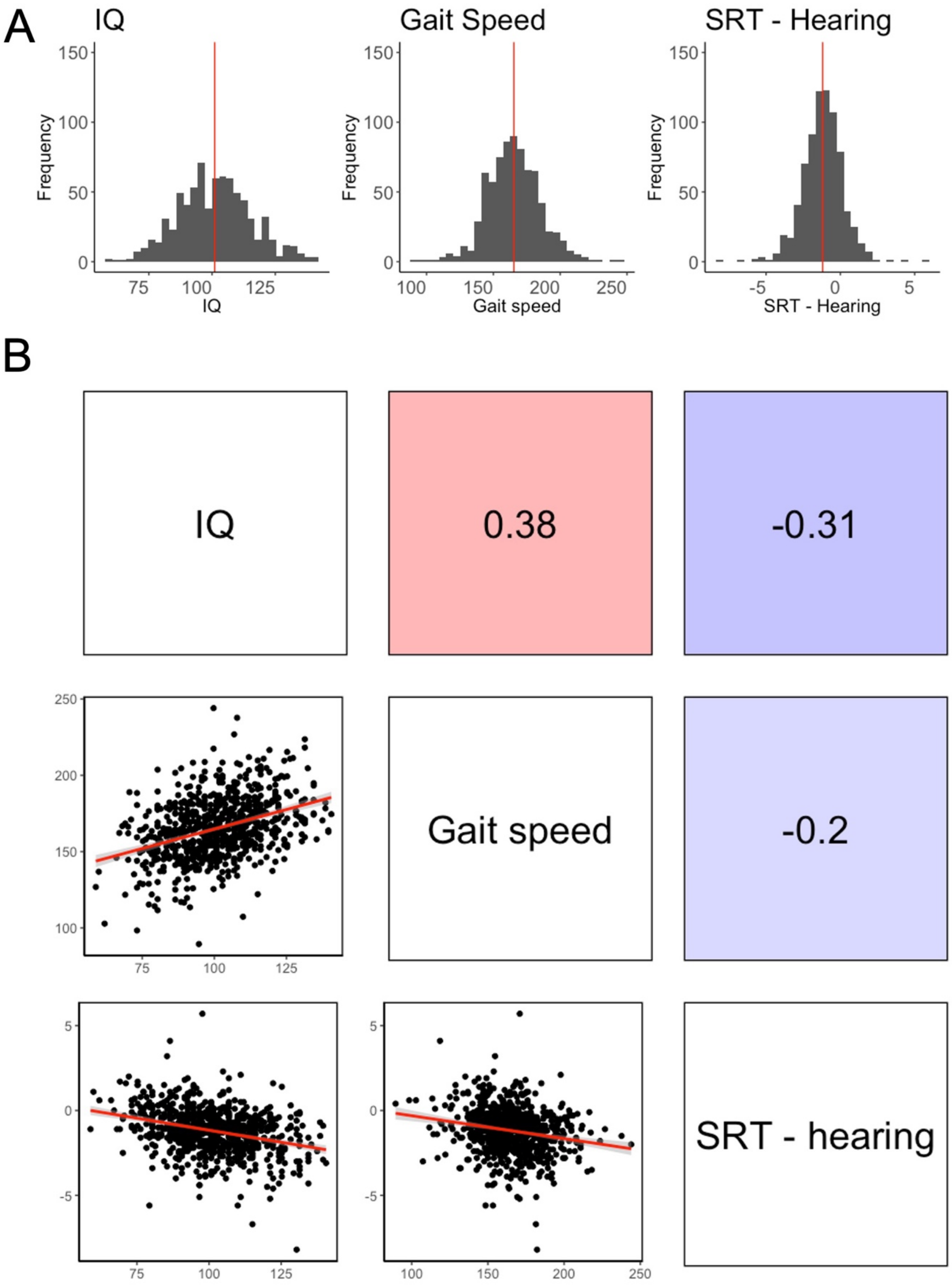

**Figure S5. Distribution and correlations of IQ, gait speed, and SRT-Hearing.** A. IQ, gait speed, and SRT-hearing all show normal distributions. Red lines represent the mean. B. Correlations between IQ, gait speed, and SRT-hearing. Upper triangle shows Pearson correlations (all *p*'s <.001) while bottom triangle shows scatterplots. Crucially, a lower SRT-hearing score indicates better hearing ability.

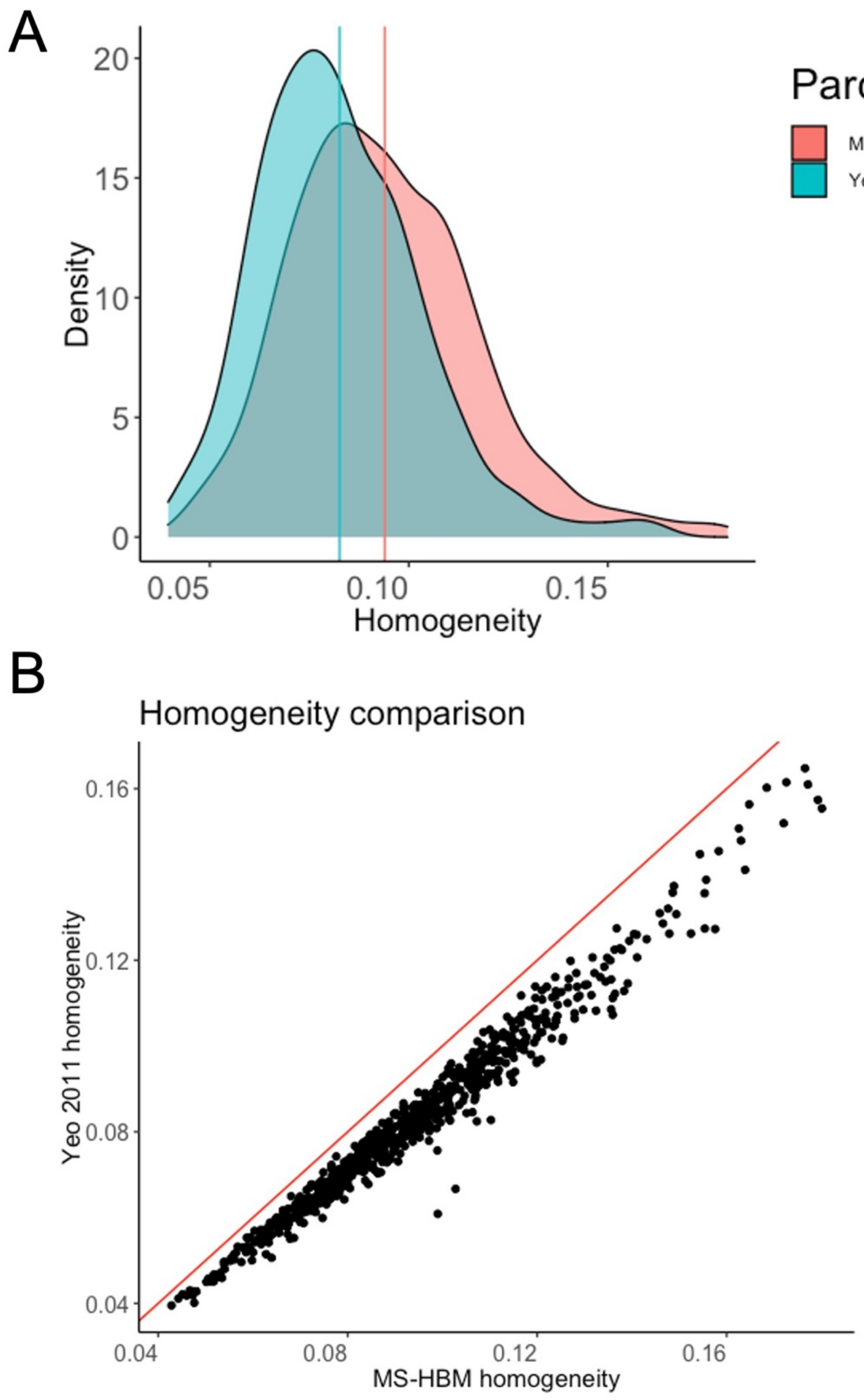

**Figure S6. Homogeneity of individualized and group parcellations.** Individualized parcellations derived using MS-HBM significantly boosted homogeneity in our sample (*t* = 10.0, *p* < .001). Pink represents homogeneity calculated from individualized parcellations and blue represents homogeneity calculated using the Yeo 2011 group average parcellation (Yeo et al., 2011). Vertical lines represent respective means.

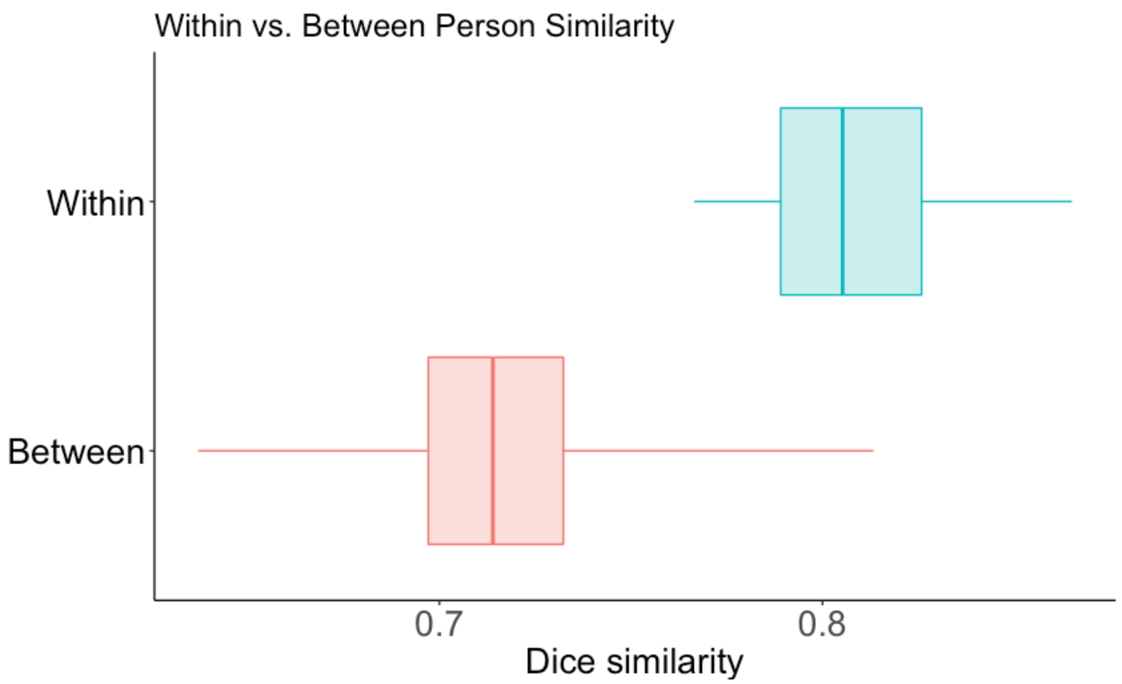

**Figure S7. Test re-test reliability of individualized parcellations.** Comparison of the within-person Dice similarity coefficients in the 19 test-retest subjects to between-person similarity. Study member's parcellations were significantly more similar to themselves at time 1 and time 2 compared to their similarity with all other test-retest Study members ($t$ = 15.1, $p$ < .001).

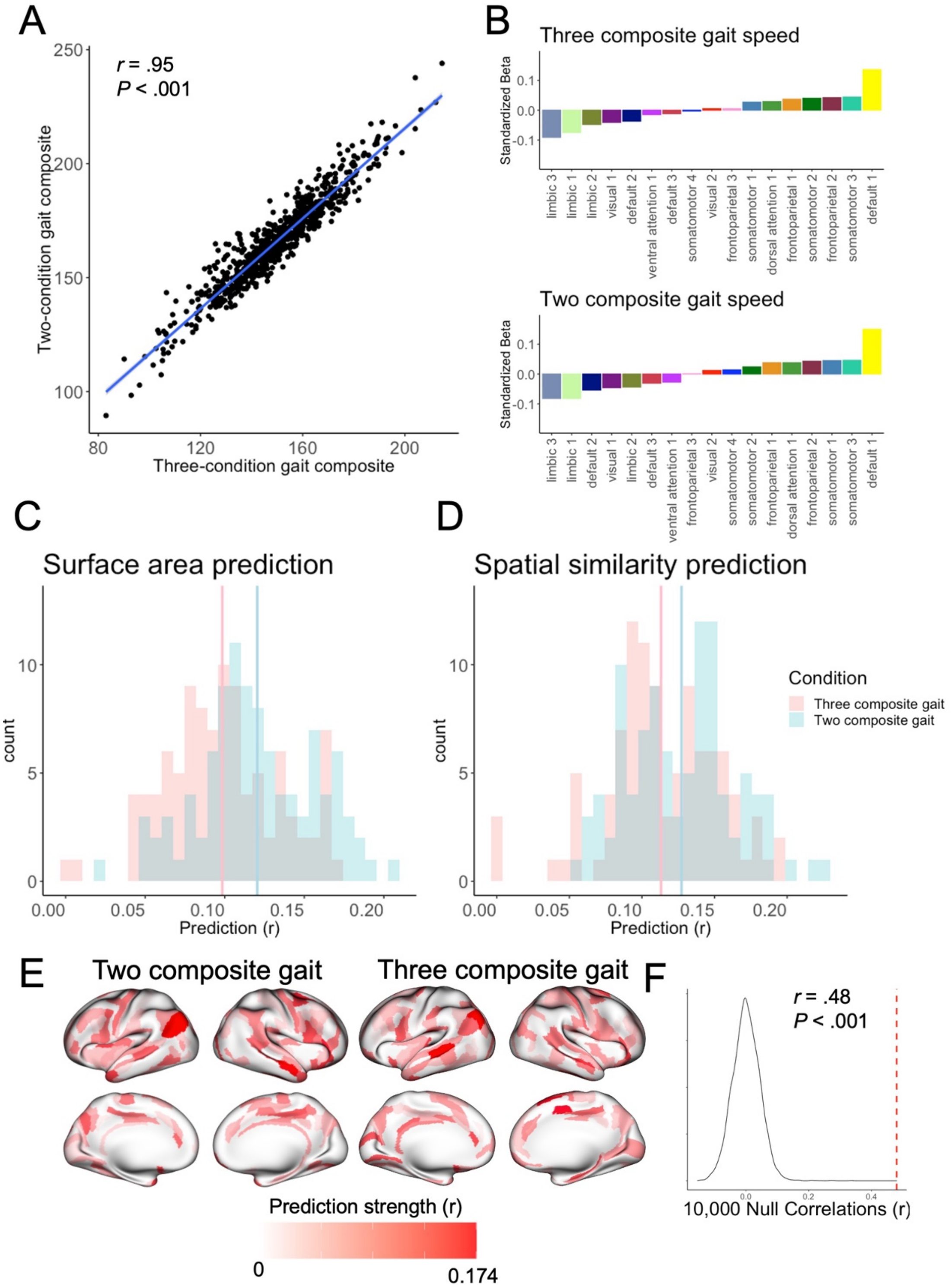

**Figure S8. Comparison of gait composites.** Two-gait composite (usual gait and maximum gait) was highly similar to the three-gait composite (usual gait, maximum gait, and dual-task gait). A. Correlation between two- and three-gait composites. B. Univariate associations between total network surface areas and each gait composite. C. Distribution of 100 random train/test split surface area predictions of two- and three-composite gait. Vertical lines represent mean prediction strength of 100 splits. C. Distribution of 100 random train/test split spatial similarity predictions of two- and three-composite gait. Vertical lines represent mean prediction strength of 100 splits. E. Regional predictions of two- and three-composite gait. F. Spin test between two- and three-composite gait regional prediction maps.

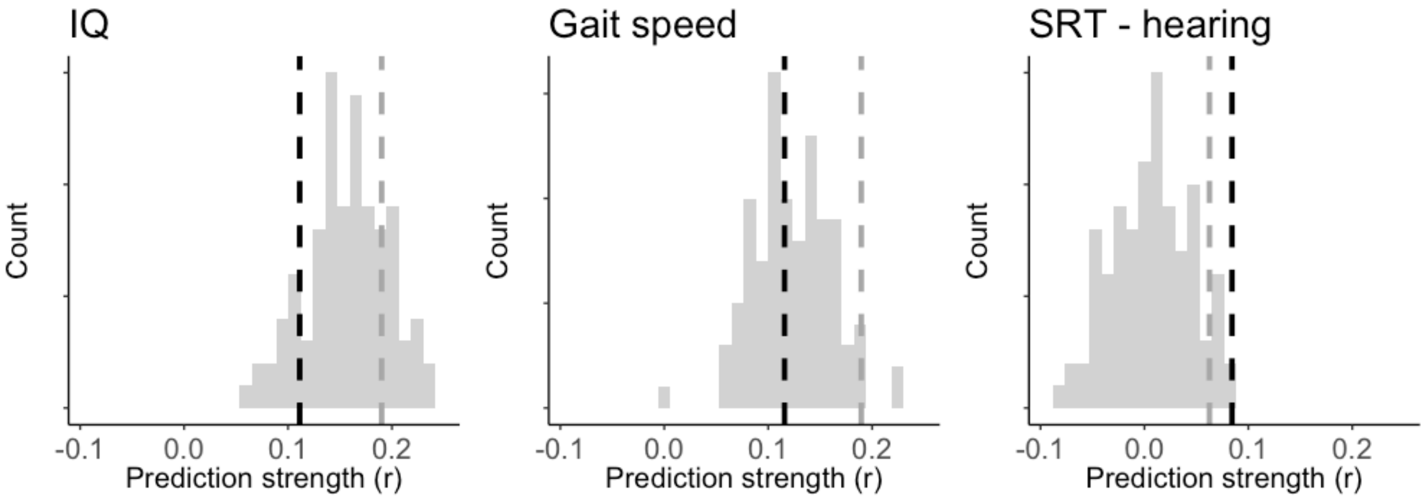

**Figure S9. Surface area prediction robustness to data splitting procedures.** Histograms show the distribution of prediction accuracies of our surface area ridge regression model based on 100 random train/test splits (each with 20 inner-loop training splits). Red vertical lines represent the prediction from our rank ordering procedure. For all three behaviors, the rank order train/test split performs comparably to random train/test splits.

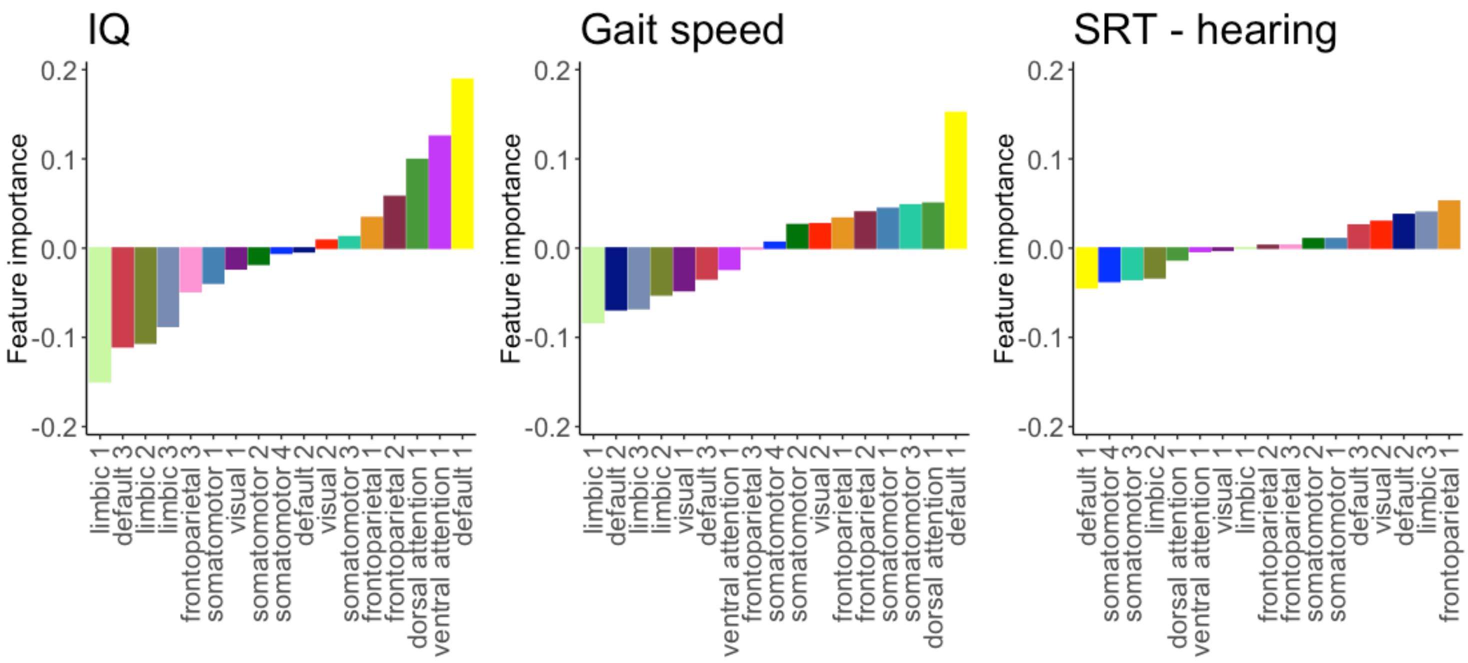

**Figure S10. Haufe transformed prediction model identifies default mode network surface area amongst the most important predictive features.** Investigation of the standardized covariates of predictor and outcome variables demonstrated that total network surface area of default mode networks was most influential in our ridge regression models trained on surface area.

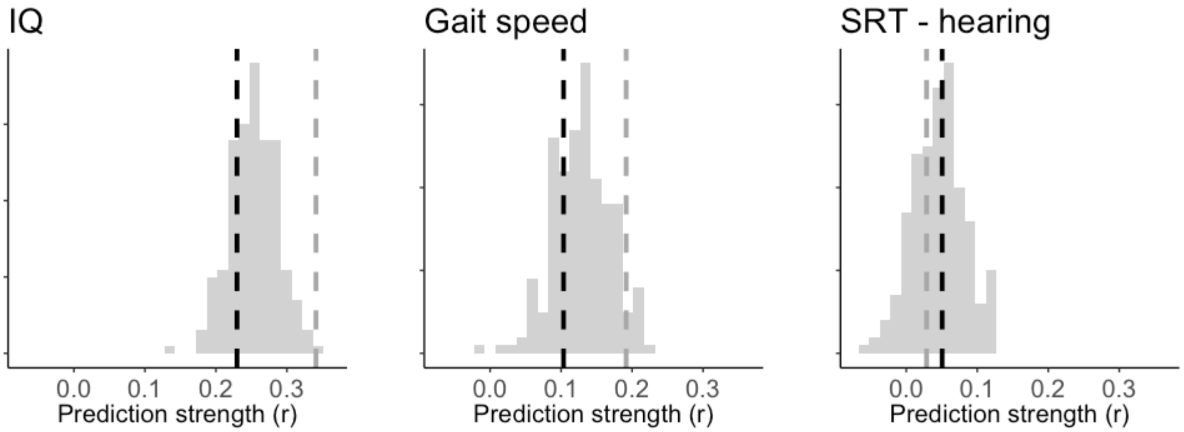

**Figure S11. Similarity prediction robustness to data splitting procedures.** Histograms show the distribution of prediction accuracies of our Dice similarity ridge regression model based on 100 random train/test splits (each with 20 inner-loop training splits). Red vertical lines represent the prediction from our rank ordering procedure. For all three behaviors, the rank order train/test split performs comparably or slightly above random train/test split

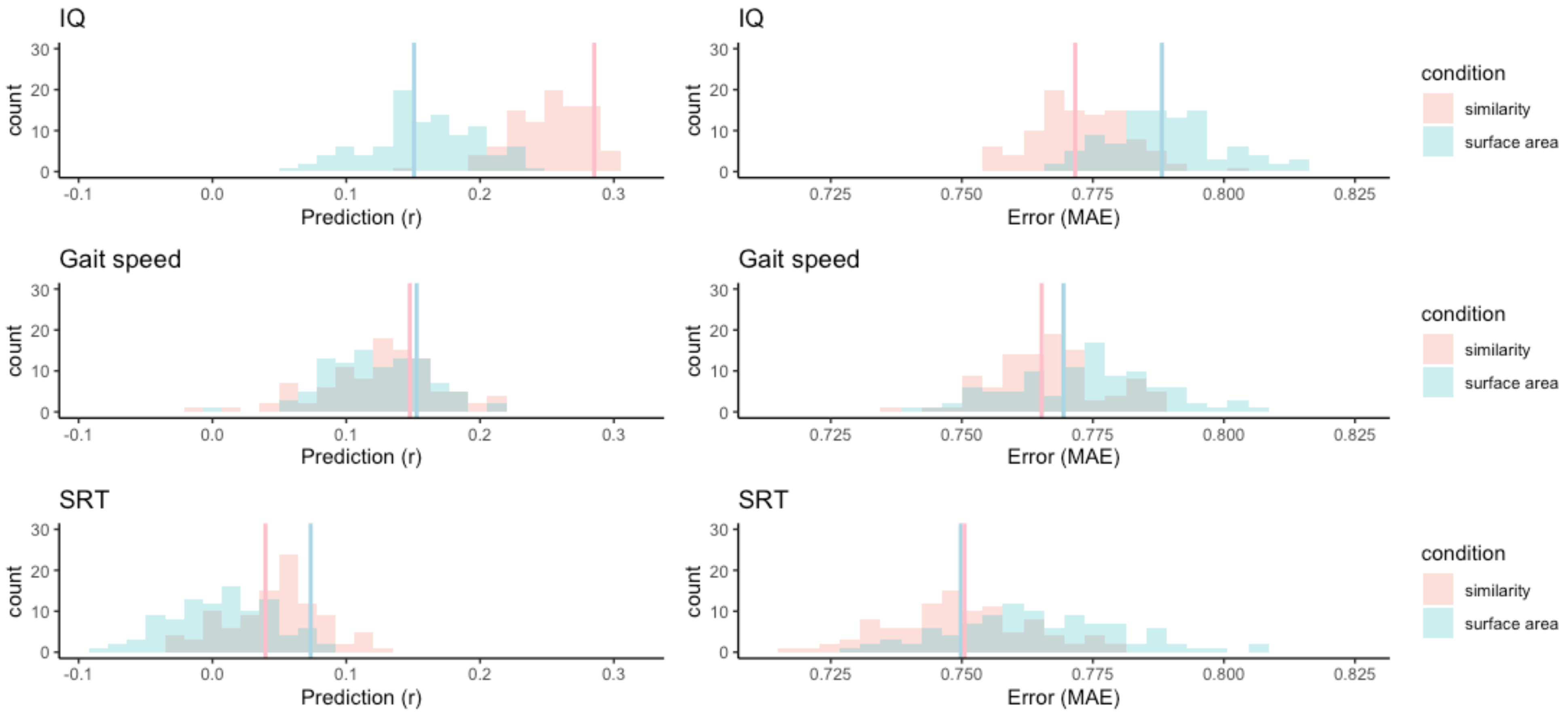

**Figure S12. Comparison of surface area and similarity predictions.** Histograms show prediction accuracy (left column) and error (right column) of 100 random train/test splits for both training methods (surface area and similarity). Vertical lines show mean prediction or mean error from both folds of our rank order data split. In general, predictions based on Dice similarity tend to have greater accuracy and less error compared to predictions based on total surface area.
